# Novel use of global occurrence data to indirectly predict suitable habitats for widely distributed marine species in data-scarce regions

**DOI:** 10.1101/2024.08.15.608095

**Authors:** Agustín M. De Wysiecki, Adam Barnett, Noela Sánchez-Carnero, Federico Cortés, Andrés C. Milessi, Gastón A. Trobbiani, Andrés J. Jaureguizar

## Abstract

This study addresses the challenge of advancing habitat use knowledge of widely distributed marine species populations when regional data is scarce. To achieve this, we use an innovative approach based on ecological niche models (ENMs) calibrated with global presence data to estimate the global niche of species, allowing for indirect predictions of suitable habitats and potential distribution in one or more regions of interest. The method leverages a range of global occurrence records, including scientific papers, government data, biodiversity repositories, and citizen science contributions, to overcome regional data scarcity, which are then integrated with environmental variables to predict habitat suitability. As a case study, we apply this method to predict suitable habitats of copper (*Carcharhinus brachyurus*) and sand tiger (*Carcharias taurus*) sharks in the Southwest Atlantic, two species of conservation concern in a region with limited data. Suitable habitats for both species were predicted, information required to guide conservation efforts. Environmental factors were key to shaping predicted distribution patterns of these large predatory sharks, aligning with previous knowledge and historical records of their latitudinal ranges. The results have significant implications for the conservation planning and sustainable management of shark populations in the Southwest Atlantic, contributing to broader efforts of marine biodiversity preservation. Additionally, the study highlights the potential of ENMs to identify essential habitats even in the absence of effort data, underscoring their value in marine conservation. This study not only advances the use of niche modelling in marine systems but also demonstrates its applicability for area-based conservation initiatives, particularly in data-poor regions.

## 1. Introduction

Ecological Niche Models (ENMs) have been used to address a wide range of theoretical and applied issues in terrestrial environments, while their applications in marine systems are relatively scarcer (Robinson et al., 2011). The main reason for this scarcity is that marine ecosystems are more dynamic and exhibit temporal variability that operate on scales ranging from hours to decades, while spatial variability can be observed on scales ranging from a few meters to thousands of kilometers (Perry & Ommer, 2003; Redfern et al., 2006). This spatiotemporal variability presents unique challenges when developing habitat models for marine species. However, over the last decade, ENMs have gained significant relevance in ecological studies in the marine realm, and their use is exponentially increasing (Robinson et al., 2017; Melo-Merino et al., 2020). The broad spatiotemporal scale of analysis achievable through the use of large databases (e.g., biodiversity repositories, citizen science) has opened up the possibility of investigating species distribution patterns across a wide range of geographical and chronological scales, which were previously difficult (or sometimes impossible) to address with traditional methods, such as field surveys. In particular, ENMs are applied in marine conservation ecology to understand the role of environmental variables in determining the distribution and habitat use of mobile species (e.g., Lucifora et al., 2015). Furthermore, ENMs also provide tools that contribute to the sustainable exploitation of marine resources, including the prediction and forecasting of species distributions and the identification of essential habitats, both of which are used to inform rational management (Valavanis et al., 2008; Bowlby et al., 2024).

The presence-only ENM is a modelling approach based solely on estimating the ecological niche of a species from presence records (Elith et al., 2006). Presence-only data consist of records describing confirmed occurrences of species, while no information is available on where they were absent, for example, sightings, species lists, biodiversity repositories, herbarium or museum specimens, etc. The use of these data presents particular challenges that need to be addressed, including errors and lack of detail in location and identification, biases in the geographic or environmental space sampled, or temporal biases in the recorded data (Newbold, 2010). Presence-only ENMs compare environmental characteristics at locations where presence records exist with those at a large number of background points and thus estimate a habitat suitability index for the species; for example, in methods such as GARP, ENFA, MaxEnt (Elith & Leathwick, 2009), or MaxLike (Royle et al., 2012). It is important to note that predictions would be more accurate if presence and absence data and/or abundance data were available, as they provide valuable information about sampling sites (i.e., enabling sample bias analysis) and prevalence (Phillips et al., 2009). When presence-absence modelling approaches (e.g., GLM, GAM) and presence-only modelling approaches using the same data sets have been compared, presence-absence models generally perform better and have higher predictive capabilities (Hirzel et al., 2001; Brotons et al., 2004). Given their characteristics, presence-only ENMs are more likely to predict potential distributions that more closely resemble the fundamental niche of the species, while presence-absence ENMs are more likely to reflect occupied areas derived from the realized niche (Zaniewski et al., 2002). However, both approaches may perform similarly when sampling effort is highly unequal (e.g., MacLeod et al., 2008). In both cases, inadequate sampling of the full range of different habitat types used by a species could lead to predictions that are not representative of reality.

Despite its inherent limitations, inference through presence-only ENMs represents a preliminary step in the development of management plans in data-scarce regions (Lopes et al., 2019). It can also provide quick and valuable information in situations of urgent conservation need. The use of data sources based on citizen activities (e.g., recreational fishing) provides the opportunity to exploit a magnitude of spatial and temporal analysis that is generally not replicable by any scientific program. For instance, it allows us to incorporate historical records covering the entire known distribution of a widely distributed species in a particular coastal region. In a broad sense, the presence-only ENM approach allows the low-cost utilization of available information based on occasional records of captures or observations from a variety of sources, in the absence of monitoring and long-term tracking programs that are difficult to implement in particular economic, social, or political contexts. Presence-only ENMs appears to be a particularly pertinent approach for identifying essential fish habitats (e.g. Owens et al., 2012; Barnett et al., 2019) for area-based conservation initiatives such as Important Shark and Ray Areas (ISRA) and implementing effective marine protected areas (Daly et al., 2018; Hyde et al., 2022; Dwyer et al., 2023).

The copper *Carcharhinus brachyurus* (Günther, 1870) and sand tiger *Carcharias taurus* Rafinesque, 1810 sharks are top benthopelagic predators found widely distributed in temperate coastal and shelf environments around the world (Compagno, 1984). These large bodied sharks exceed 3 meters in length, exhibit slow growth, late maturation, and long lifespans (Gilmore et al., 1983; Walter & Ebert, 1991). Both species face significant threats from industrial, small-scale, and recreational fisheries, where they are both targeted and caught as bycatch using various gear types, mainly demersal longline and gillnet and to a lesser extent, pelagic longline and demersal trawl (e.g., Chiaramonte, 1998; Dudley & Cliff, 2010). Both species have experienced substantial population declines, suspected to be around 30-50% for *C. brachyurus* and over 80% for *C. taurus* (Huveneers et al., 2020; Rigby et al., 2021). As a result, *C. brachyurus* is assessed as Vulnerable by the IUCN, while *C. taurus* is classified as Critically Endangered (Huveneers et al., 2020; Rigby et al., 2021), highlighting the need for ongoing conservation efforts for both species. The wide distribution and possible large-scale movements of both shark species implies that special use incorporates habitats across multiple jurisdictions and/or national boundaries throughout their lives (Huveneers et al., 2021; Rogers et al., 2022; Dwyer et al., 2023). This exposes them to a multitude of factors that directly affect their conservation status, including fishing, pollution, habitat loss, and climate change (Jaureguiberry et al., 2022). Consequently, the implementation and effectiveness of management measures are rendered more complex (Campana, 2016). In particular, *C. taurus* has been classified as an Endangered migratory species under the Convention on the Conservation of Migratory Species of Wild Animals (CMS, 2024), which reinforces the necessity to develop regional, cross jurisdictional and global studies to derive spatial management tools with the aim of improving the protection of essential habitats and the migratory routes that connect them.

There are at least four regions worldwide where populations of both species coexist and stocks appear to be stable, including the Southwest Atlantic, Mediterranean Sea, Southern Africa, and Australia (Figure S1.1). In other coastal areas, while the species do not overlap, they are abundant individually, such as *C. brachyurus* in New Zealand (Francis, 1998; Kellett, 2021), and *C. taurus* along the east coast of the United States (Branstetter & Musick, 1994; Teter et al., 2014). Despite evident connectivity within each region, genetic studies indicate phylogeographic discontinuity and reduced overall connectivity between different populations of each species, particularly between hemispheres for *C. taurus* (Ahonen et al., 2009; Lucifora et al., 2003; Stow et al., 2006), with Oceania being an exception for *C. brachyurus* (Benavides et al., 2011; Junge et al., 2019). In the Southwest Atlantic (SWA), literature indicates that *C. brachyurus* is distributed from Guarapari, in the state of Espírito Santo, Brazil (northernmost record at 20°41’S) (Cuevas et al., 2022), to the San Matías Gulf, in the province of Río Negro, Argentina (southernmost record at ∼41°S) (Perier et al., 2011). Similarly, the distribution of *C. taurus* extends from Itanhaém, in the state of São Paulo, Brazil (northernmost record at 24°11’S) (Motta, 2006), to the San Matías Gulf, in the province of Río Negro, Argentina (southernmost record at 41°05’S) (Menni et al., 1981). Both species represent significant fishing resources that have been and continue to be targeted by recreational fishing since the 1950s (Cedrola et al., 2011). In particular, *C. brachyurus* has also been target of small-scale artisanal fishing (Chiaramonte, 1998a, 1998b). Additionally, these two species are incidentally caught in various artisanal and industrial fisheries in the region (e.g. Marín et al., 1998; Jaureguizar et al., 2015; Mas et al., 2014; Silveira et al., 2018). It is important to note that fisheries have been dynamic and have changed over time. Some fishing methods that were common in the past may not be applicable today due to changes in fishing gear or for economic reasons that have led fleets to move to other fishing areas. However, given the fishing exploitation, the relative abundances of their populations in the long term show a strong decrease (Barbini et al., 2015; Irigoyen & Trobbiani, 2016). As a result, *C. brachyurus* and *C. taurus* are included in national and regional action plans for their conservation and management, including the PAN-Tiburones Uruguay (Domingo et al., 2008), PAN-Tiburones Argentina (CFP, 2009), PAN-Tubarões Brasil (Kotas et al., 2023), and PAR-Tiburones (CTMFM, 2018). Although some biological information (abundance, distribution, feeding, growth, and reproduction) has been documented during the last two decades (Table S1.1), knowledge about large-scale habitat use and distribution of these species along the SWA remains limited.

In contrast to other predatory sharks commonly caught across various fisheries and field surveys in the SWA, such as the school (*Galeorhinus galeus*) and sevengill (*Notorynchus cepedianus*) sharks (e.g., Chiaramonte et al., 2016; De Wysiecki et al., 2018), data on the occurrences of *C. brachyurus* and *C. taurus* are sparse and do not adequately depict their distribution in the study region (Figure S1.2). Previous studies assumed that the spatial distribution of data collection for *G. galeus* and *N. cepedianus* sufficed to model their regional niches (De Wysiecki et al., 2020, 2022b). Conversely, the available data for *C. brachyurus* and *C. taurus* suggest an insufficient spatial representation of known occurrences, posing an initial obstacle in predicting their potential distribution in the region. The goal of this study is to predict the potential distribution of *C. brachyurus* and *C. taurus* populations in the SWA under regional data-poor conditions. This is achieved through an alternative, yet conceptually similar, innovative ENM approach. The method involves calibrating and modelling the ecological niche of these species using global data (i.e., considering the rest of the world’s populations) and then transferring the habitat suitability index to the SWA, thereby facilitating the indirect mapping of their potential distribution. The underlying assumption of this analytical approach is that the world’s populations of a species exhibit similar habitat use patterns. This hypothesis was recently tested using global occurrences of *N. cepedianus*, demonstrating that the global model effectively identified the species’ core distribution (De Wysiecki et al., 2022). Additionally, the model successfully predicted occupied areas at the extremes of *N. cepedianus*’ distribution before field surveys confirmed their presence (Schulte et al., 2024).

## 2. Materials and Methods

The goal of the ENM approach was to predict (continuous and binary) annually averaged (annual model) suitable areas of the SWA for two species, the copper (*Carcharhinus brachyurus*) and sand tiger (*Carcharias taurus*) sharks. To promote transparency and reproducibility, we described the methodological approach following the Overview, Data, Model, Assessment, Prediction (ODMAP) protocol for species distribution models (Zurell et al., 2020). In the following sections, we provide an overview of methods; a detailed description of the five ODMAP components can be found in Supplement 2. All analyses were developed with R version 4.2.2 (R Core Team, 2022), and the code is available on GitHub (Agustindewy/Copper_Stiger_shark_SWA).

### 2.1 Occurrence records

The occurrence records of *C. brachyurus* and *C. taurus* were collected from various data sources to maximize the number of records in the SWA (Table 1). Open-access records were obtained from published literature, including indexed and grey articles, biodiversity repositories, and social media. An exhaustive literature review was conducted using Google Scholar with the terms ‘Carcharhinus brachyurus’ and ‘Carcharias taurus’ to identify articles reporting the species’ catches in the SWA. When only the catch location was reported, the occurrence record was manually georeferenced. Additionally, all *C. brachyurus* and *C. taurus* records were downloaded both from the Global Biodiversity Information Facility (GBIF, 2021a; 2021b) and Ocean Biodiversity Information System (OBIS, 2021). Social media records were primarily sourced from Facebook posts within public angler groups (refer to ODMAP for details). Additional records from Argentina were requested, including commercial logbook data from Pesca Nación and scientific surveys from the Instituto Nacional de Investigación y Desarrollo Pesquero (INIDEP).

**Table 1.** Brief description of the data sources consulted for occurrence records of *Carcharhinus brachyurus* (CB) and *Carcharias taurus* (CT). The records are divided between the Southwest Atlantic (SWA) region and the rest of the world (Global). The number of occurrences collected from each source is indicated. Data collection was concluded on February 28, 2022.

| Source | Brief description | SWA |  | Global |  |
| --- | --- | --- | --- | --- | --- |
|  |  | CB | CT | CB | CT |
| Publications | Articles in indexed journals | 21 | 61 | 44 | 114 |
| Grey literature | Institutional reports, fishing magazines, and theses | 156 | 19 | 0 | 0 |
| Biodiversity repositories | GBIF and OBIS | 2 | 2 | 10769 | 2323 |
| Government data | Observer/logbook data and surveys | 59 | 16 | 1475 | 22 |
| Professional contact | Surveys | 74 | 1 | 0 | 0 |
| Social media | Facebook and YouTube | 104 | 21 | 0 | 0 |
| Total |  | 416 | 120 | 12288 | 2459 |

GBIF = Global Biodiversity Information Facility, OBIS = Ocean Biodiversity Information System

The search for *C. brachyurus* and *C. taurus* occurrence data was expanded globally (Table 1). In addition to the scarce records compiled for these two species in the SWA, global occurrences from freely accessible publications indexed in Google Scholar, and GBIF and OBIS repositories were included. Records from national observation programs aboard commercial fishing vessels in Australia (*C. brachyurus* and *C. taurus*) and New Zealand (*C. brachyurus*) were also requested. These data are subject to current confidentiality agreements and were provided by the Australian Fisheries Management Authority (AFMA) and Fisheries New Zealand (FNZ).

### 2.2 Environmental predictors

To delineate the ecological niche of *C. brachyurus* and *C. taurus*, a comprehensive suite of environmental predictors with global coverage was considered. These predictors include distance to the coast, depth, bottom temperature, bottom salinity, surface temperature, Kd490 coefficient, primary productivity, and the magnitude of thermal fronts. The selected predictors have direct or indirect effects, e.g. through prey availability (Barnett & Semmens, 2012), on the movement and habitat use of sharks and rays (Schlaff et al., 2014), and are widely used in both niche modelling (Bradie & Leung, 2017) and elasmobranch remote sensing studies (Williamson et al., 2019). In particular, the distance to the coast aims to delineate significant coastal habitats (e.g., spawning, breeding, and feeding areas), while depth enables the description of topography in light of their demersal habits. Temperature, salinity, the diffuse attenuation coefficient Kd490, primary productivity, and the magnitude of thermal fronts were all considered to characterize the physical conditions of the water column. Temperature and salinity serve as indicators of the species’ physiological limits (Schlaff et al., 2014). The Kd490 coefficient reflects the amount of sediment dissolved in turbid water, which is interpreted as a possible source of protection and refuge for species, particularly juveniles (Chin et al., 2013; Jaureguizar et al., 2023). Finally, primary productivity and the magnitude of thermal fronts, derived from surface temperature images following the methodology proposed by (Scales et al., 2014a), underscore the significance of carbon and temperature gradients as markers of possible large-scale food sources (Alemany et al., 2009; Scales et al., 2014b).

The selected environmental predictors are assumed to be sufficient to characterize and capture the environmental conditions that allow the species to maintain their populations over time. The characteristics of the environmental predictors, including source, spatial resolution, range of values, units, and analysis period, are detailed in Table 2. The oceanographic variables, including bathymetry, were downloaded from the global Bio-ORACLE marine dataset (version 2.1), at a resolution of 5 arcmins (∼0.083°) and a global extent (Assis et al., 2018). The distance to the coast was downloaded from the Global Self-consistent, Hierarchical, High-resolution Geography Database (version 2.3.7), at a resolution of 1 arcmins (∼0.016°) and a global extent (Wessel & Smith, 1996). A spatial resolution of 5 arcmins was determined for modelling, as it was the lowest resolution among the predictors. Predictors with higher native resolutions were resampled, possible data gaps were filled through neighbouring cell interpolation, and different rasters were clipped by the coastline polygon (refer to ODMAP for details).

**Table 2.** Description of the environmental predictors used to model the global ecological niche of *Carcharhinus brachyurus* and *Carcharias taurus* shark species.

| Predictors | Source | Native resolution | Range (min – max) | Units | Period |
| --- | --- | --- | --- | --- | --- |
| Distance to coast | MARSPEC | 0.016° | 1 – 2777.8 | km | N/A |
| Depth | Bio-ORACLE | 0.083° | -10494 – 0 | m | N/A |
| Bottom temperature | Bio-ORACLE | 0.083° | -1.8 – 31.4 | °C | 2000 – 2014 |
| Bottom salinity | Bio-ORACLE | 0.083° | 5 – 40.8 | PSU | 2000 – 2014 |
| Surface temperature (SST) | Bio-ORACLE | 0.083° | -1.8 – 32.9 | °C | 2000 – 2014 |
| Thermal fronts | SST derived | 0.083° | 0 – 1.77 | °C | 2000 – 2014 |
| Kd490 diffuse attenuation | Bio-ORACLE | 0.083° | 0.01 – 0.97 | m<sup>-1</sup> | 2000 – 2014 |
| Primary productivity | Bio-ORACLE | 0.083° | 0 – 0.38 | g C m<sup>-2</sup> day<sup>-1</sup> | 2000 – 2014 |

MARSPEC = Ocean Climate Layers for Marine Spatial Ecology, Bio-ORACLE = Marine data layers for ecological modelling

### 2.3 In-situ vs modelled depth discrepancies

The most representative data of *C. brachyurus* captures were mainly collected in Australia (*n* = 9995) and New Zealand (*n* = 291), using various bottom fishing gears such as longlines, trawl nets, and gillnets. Through on-site measurements, these captures revealed consistent patterns regarding the depths at which the species was caught. In Australia, captures were typically (99th percentile) recorded at depths shallower than 450 m, with a maximum depth of 611 m. In New Zealand, usual depths were below 578 m, reaching a maximum depth of 731 m. In other parts of the world (*n* = 60), *C. brachyurus* was captured at average depths below 364 m, with a maximum depth of 536 m. However, when comparing these measurements with data provided by the bathymetric layer, very high values were observed, up to 4361 m (Figure S1.3). At least 3% of the records (12.4% if duplicate captures in the same space are excluded) presented depth values exceeding the maximum recorded depth of 731 m.

Similarly, the most representative data of *C. taurus* captures were primarily collected in Australia (*n* = 183) and the Northwest Atlantic (*n* = 310), using various bottom fishing gears. In Australia, captures were typically (99th percentile) recorded at depths shallower than 555 m, with a maximum depth of 640 m. In the Northwest Atlantic, usual depths were below 448 m, reaching a maximum depth of 700 m. In other parts of the world (*n* = 41), *C. taurus* was captured at average depths below 78 m, with a maximum depth of 90 m. However, when comparing these measurements with data provided by the bathymetric layer, very high values were observed, up to 5081 m (Figure S1.4). At least 1.7% of the records (5.9% if duplicate captures in the same space are excluded) presented depth values exceeding the maximum recorded depth of 700 m.

This discrepancy between field-measured depths and those derived from bathymetric models is due to measurement and estimation errors occurring on steep slopes and abrupt drops along the continental shelf margin (De Wysiecki et al., 2022). As a result, depth was discarded as a possible predictor of ecological niche, and the distance to the coast variable was retained to avoid estimating excessively wide environmental ranges for the species. This issue stems from spatial resolution and could similarly affect other variables for which no in situ data is available and cannot be verified. However, this was not expected to occur because their distributions are more gradual or smoother, as is the case with the distance to the coast. Consequently, seven final predictors (distance to the coast, bottom temperature, bottom salinity, surface temperature, Kd490 coefficient, primary productivity, and the magnitude of thermal fronts) were considered to estimate the ecological niche of the global population of *C. brachyurus* and *C. taurus*.

### 2.4 Calibration and evaluation

The analyses consisted of an annual model for the populations of *C. brachyurus* and *C. taurus* on a global scale. These models consider the total occurrences of the species worldwide and averaged environmental values with global coverage. To minimize the spatial clustering of calibration sets, we utilized a thinning distance of 50 km with the “spThin” R package (Aiello-Lammens et al., 2015). To reduce the effects of sampling bias and clustering in environmental space, we applied an environmental filter using the “envSample” custom function. This filter used thresholds of 0.5°C for surface temperature and 1 km for distance to the coast (Varela et al., 2014). Accessible areas were then defined using 1000 km buffer polygons around each calibration record to determine the calibration area (i.e., M space, Soberón & Peterson, 2005). This buffering distance was chosen based on tracking data, which shows that both species are capable of movements exceeding 1000 km. In southern Australia and the Southwest Atlantic, *C. brachyurus* exhibited movements of up to ∼2500 km between tagging and recapture sites (Rogers et al., 2013; Cuevas et al., 2022). On the east coast of Australia, unidirectional movements of *C. taurus* of up to ∼1200 km were recorded in coastal areas (Bansemer & Bennett, 2011). However, these large distances occurred within the central distribution area of each species. Therefore, polygons with a diameter of 1000 km were used, as it is unlikely that larger polygons would adequately represent the accessible areas along the boundaries of their distributions. The calibration area was used to randomly select background data (*n* = 10,000) for model calibration.

To map the potential distribution of species, global niche models require the transfer of the habitat suitability index to areas of the world where representative data are lacking (e.g., SWA) and lie outside the model’s calibration area. The process of transferring the niche to new areas can be complicated if there are new environmental combinations not covered in the model, resulting in extrapolation in estimates (Qiao et al., 2019). Therefore, it is crucial to assess the models’ ability to extrapolate to new areas (Wenger & Olden, 2012; Yates et al., 2018). One way to select models with better extrapolation capability is by using spatial blocks as a method of data partitioning during cross-validation, ensuring comparison between environmental combinations from different geographic areas (Muscarella et al., 2014). To achieve this, an automated calibration and evaluation protocol was implemented using the ENMevaluate function from the R package "ENMeval" to compare candidate models of different complexities and provide flexibility in creating spatial blocks during evaluation (Kass et al., 2021). In this protocol, both calibration data and background points are divided into spatial blocks (k = 5) for cross-validation. During this evaluation process, one block is used for calibration and the remainder (k - 1 blocks) for iteratively evaluating model fit until all blocks have been used. To significantly reduce spatial autocorrelation in the data, the effective block size over which observations are spatially independent was determined (Valavi et al., 2019). The effective size and spatial blocks were determined using the spatialAutoRange and spatialBlock functions from the R package "blockCV," respectively (Valavi et al., 2019).

The models were calibrated using version 3.4.3 of the MaxEnt algorithm. Since MaxEnt’s default settings do not always lead to the best model performance (Shcheglovitova & Anderson, 2013; Radosavljevic & Anderson, 2014), subsets of entity classes producing simpler linear and unimodal fits were considered (i.e., linear + quadratic, linear + product, and linear + quadratic + product), along with a complete set of regularization multipliers (i.e., a sequence from 0.1 to 2 in increments of 0.1, plus 2.5, 3, 4, 5, 7, and 10). The automated evaluation protocol implemented in the R package "kuenm" was then applied to compare candidate models of different complexity (Cobos et al., 2019). The protocol selects the best models through three steps. Firstly, it identifies the most significant models using the partial area under the receiver operating characteristic curve (pROC) (Peterson et al., 2008), which has been shown to be a more appropriate indicator than the total area under the ROC curve (Lobo et al., 2008; Jiménez-Valverde, 2012). Secondly, it selects the best-performing models using omission rates (5% error threshold), which represent the proportion of cells with species presence falling outside the predicted suitable areas for the species. For this, presence data are randomly divided into 75% for calibration and 25% for internal cross-validation. The omission rate indicates how well models built with calibration data predict the occurrence of evaluation data (Anderson et al., 2003). Thirdly, it selects the simplest models according to the corrected Akaike information criterion for small sample sizes (AICc) (Warren & Seifert, 2011). The AICc values indicate how well the models fit the data while penalizing complexity and favoring simpler models (selection criterion ΔAICc ≤ 2). Finally, response curves were constructed for each final model to understand how habitat suitability changes as the values of each predictor vary, while keeping the rest of the predictors at their average value. The curves were created using the R package "SDMtune" (Vignali et al., 2020).

### 2.5 Global prediction and risk of extrapolation

The final niche model was transferred worldwide, excluding the Arctic and Antarctic territories. This allowed for the prediction of extensive regions potentially suitable for *C. brachyurus* and *C. taurus* in geographic space. Areas of strict extrapolation, represented by environmental combinations entirely outside those considered during model calibration, were excluded from prediction through Mobility-Oriented Parity analysis (MOP) (Owens et al., 2013).

## 3. Results

### 3.1 Carcharhinus brachyurus

The annual model prediction, based on the fitting of the global population of *C. brachyurus*, revealed extensive areas of suitable habitat in temperate regions of both hemispheres (Table 3, Figure 1a). These regions include the Northeast, Southeast, and Northwest Pacific, Northwest, Northeast, Southwest, and Southeast Atlantic, as well as Australia and New Zealand. When comparing these results with the species’ presence distribution in space, it is observed that the suitable regions of the Northwest Atlantic and the Azores Islands represent invadable areas for the species that have not yet been occupied (Figure 1ab). Regarding the predictors used in the global population model, it was found that bottom temperature contributed most significantly (41.1%), followed by distance to the coast (38.6%), and surface temperature (12.9%) (Table 3). Response plots indicated that the ecological niche of the global population of *C. brachyurus* is mainly associated with coastal waters (<100 m depth), with optimal temperatures both at the bottom (range of 9–26°C) and at the surface (range of 14–23°C) (Figure 1c). Additionally, a marginal association was also observed with marine waters (>32 PSU), slightly turbid (optimal range of 0.05–0.37 m^-1^), and with mild levels of productivity (optimal range of 0.002–0.043 gCm^-2^ day^-1^) and thermal fronts (>0.08°C) (Figure S1.5).

**Figure 1.**
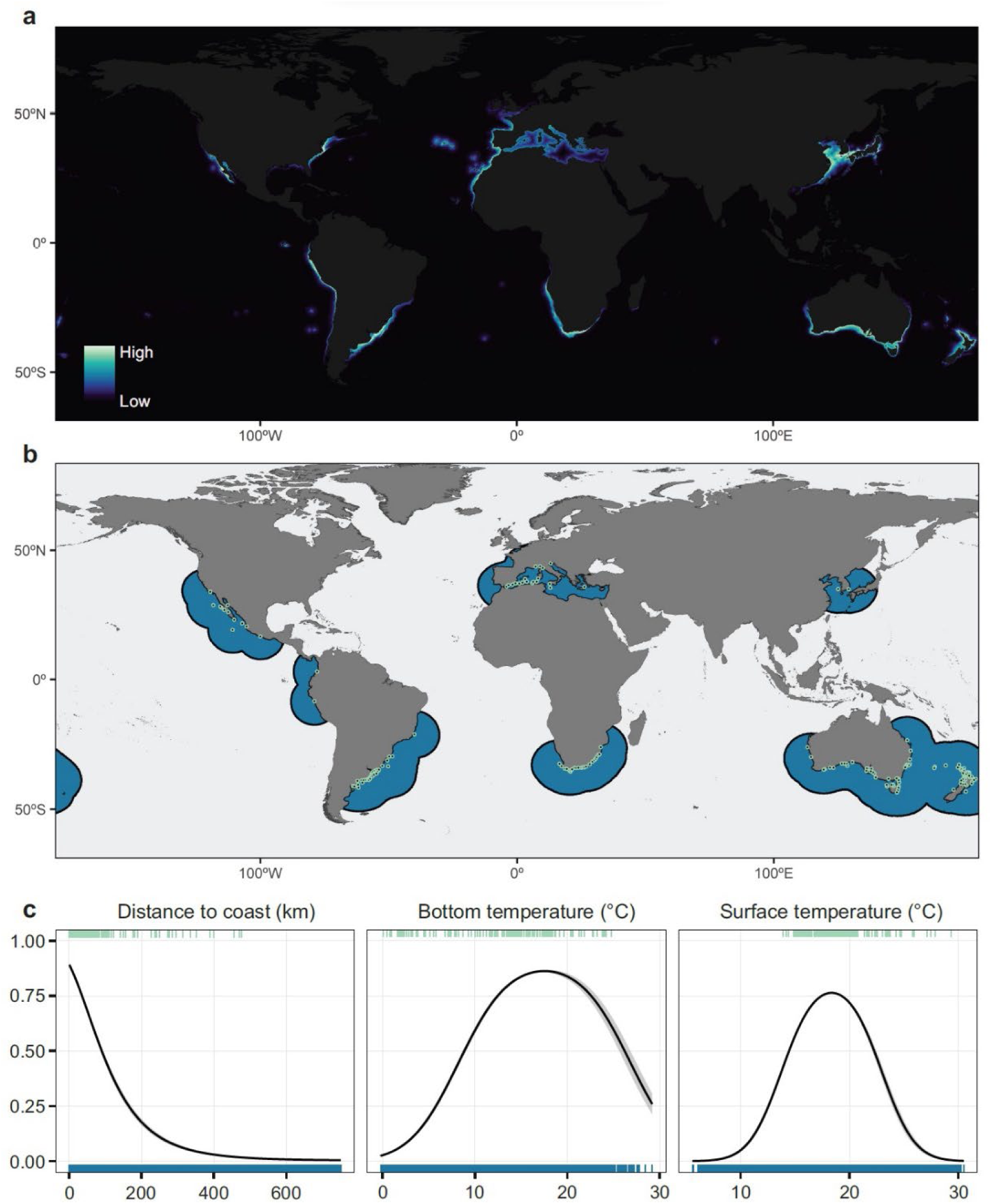
Main model results for the global population of *Carcharhinus brachyurus*, including: (**a**) Continuous prediction of habitat suitability representing the potential distribution of the species worldwide, covering both occupied and potentially invadable areas. Areas of strict extrapolation due to model transfer were removed. (**b**) Calibration areas constructed as 1000 km diameter buffer circles around each of the calibration points, represented as circles. (**c**) Response plots (logistic output) of the most influential predictors in the model. The curves illustrate how the prediction varies as the values of each predictor are modified, while keeping the rest of the predictions at their average value. The lines represent the mean response, and the shaded area corresponds to one standard deviation of variability from ten MaxEnt replicates. At the top, predictive values corresponding to calibration points are shown, while at the bottom, values corresponding to 10,000 background locations randomly taken from the calibration area are displayed.

**Table 3.**
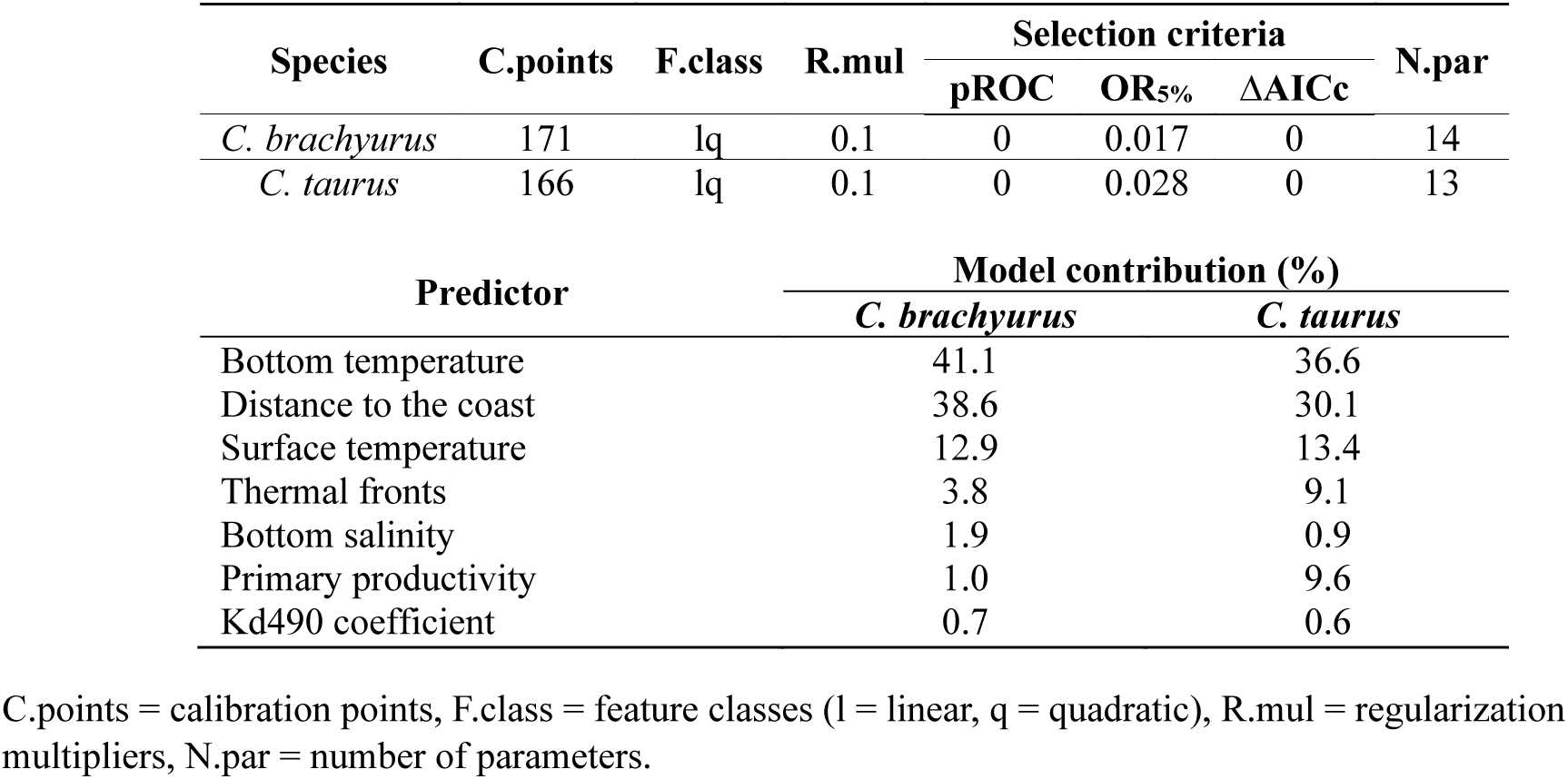
Parameterization of the best annual models selected for the global population of *Carcharhinus brachyurus* and *Carcharias taurus*. The models were selected using three criteria: the partial area under the receiver operating characteristic curve (pROC), the 5% omission rate (OR_5%_), and the Akaike information criterion corrected for sample size (ΔAICc < 2). Predictors were ranked in descending order based on their contribution to the model.

The model predicts that suitable habitat for *C. brachyurus* is widely distributed in the area of interest, the SWA (Figure 2a). The areas of potential distribution were found in coastal and continental shelf zones, spanning from southern Brazil (∼22°S) to central Argentina (∼44°S) (Figure 2b). Estuarine areas such as the Río de la Plata and Lagoa dos Patos, areas beyond the continental slope, and those south of the parallel 44°S were not suitable for the species. Notably, the overlap of historical occurrences of *C. brachyurus* in the region showed a large number of points in areas away from the coast that were predicted as unsuitable for the species (Figure 2a). When comparing the environmental combinations occupied by these occurrences with those occupied by the calibration points of the global-scale model, it was observed that the occurrences occupied habitats significantly farther from the coast (Figure S1.6).

**Figure 2.**
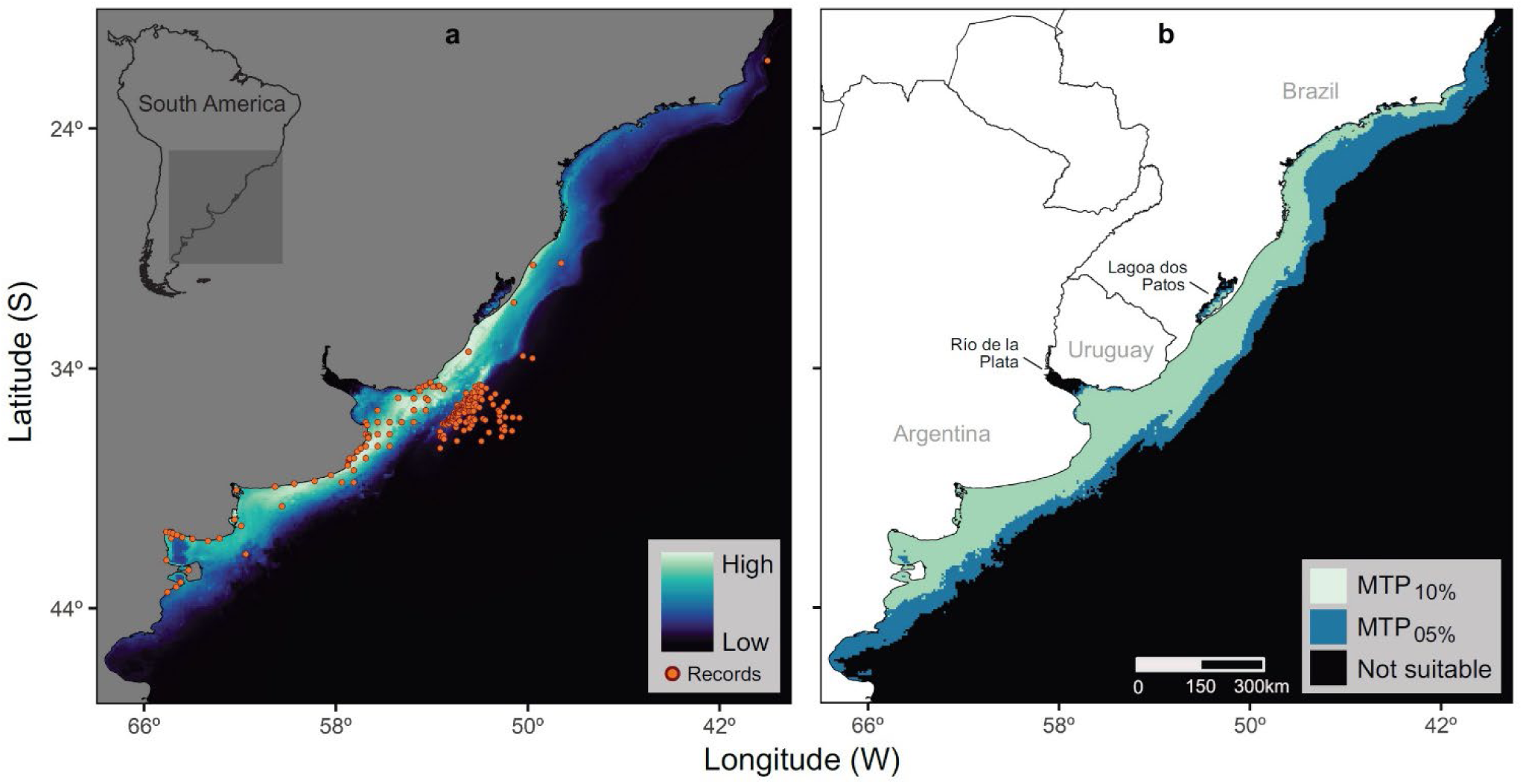
Continuous (**a**) and binary (**b**) potential distribution of the *Carcharhinus brachyurus* population in the Southwest Atlantic, based on the indirect estimation of its ecological niche at a global scale. The predictions are not limited to the calibration area but represent the model transfer to the entire region, while areas of strict extrapolation have been removed. Historical occurrences of the species in the region are shown in **a**. The binary prediction was established using minimum training presence (MTP) thresholds with 10% and 5% error on the habitat suitability prediction.

### 3.2 Carcharias taurus

The annual model prediction, based on the fitting of the global population of *C. taurus*, revealed extensive areas of suitable habitat in subtropical and temperate regions of both hemispheres (Table 3, Figure 3a). These regions include the Northeast, Southeast, and Northwest Pacific, Northwest, Northeast, Southwest, and Southeast Atlantic, as well as Australia and New Zealand. When comparing these results with the species’ presence distribution in space, it is observed that the suitable regions of the East Pacific, Galápagos Islands, Azores Islands, Madagascar, and New Zealand represent invadable areas for the species that have not yet been occupied (Figure 3ab). The predictor that contributed the most to the model was bottom temperature (36.6%), followed by distance to the coast (30.1%), and surface temperature (13.4%) (Table 3). Response plots indicated that the ecological niche of the global population of *C. taurus* was mainly associated with coastal waters (<100 m depth), with subtropical and temperate temperatures, both at the bottom (optimal range of 10–30°C) and at the surface (optimal range of 16–27°C) (Figure 3c). Additionally, a marginal association was also observed with marine waters (>33 PSU), slightly turbid (optimal range of 0.08–0.45 m^-1^), and with mild levels of productivity (optimal range of 0.005–0.045 gCm^-2^ day^-1^) and thermal fronts (>0.1°C) (Figure S1.7).

**Figure 3.**
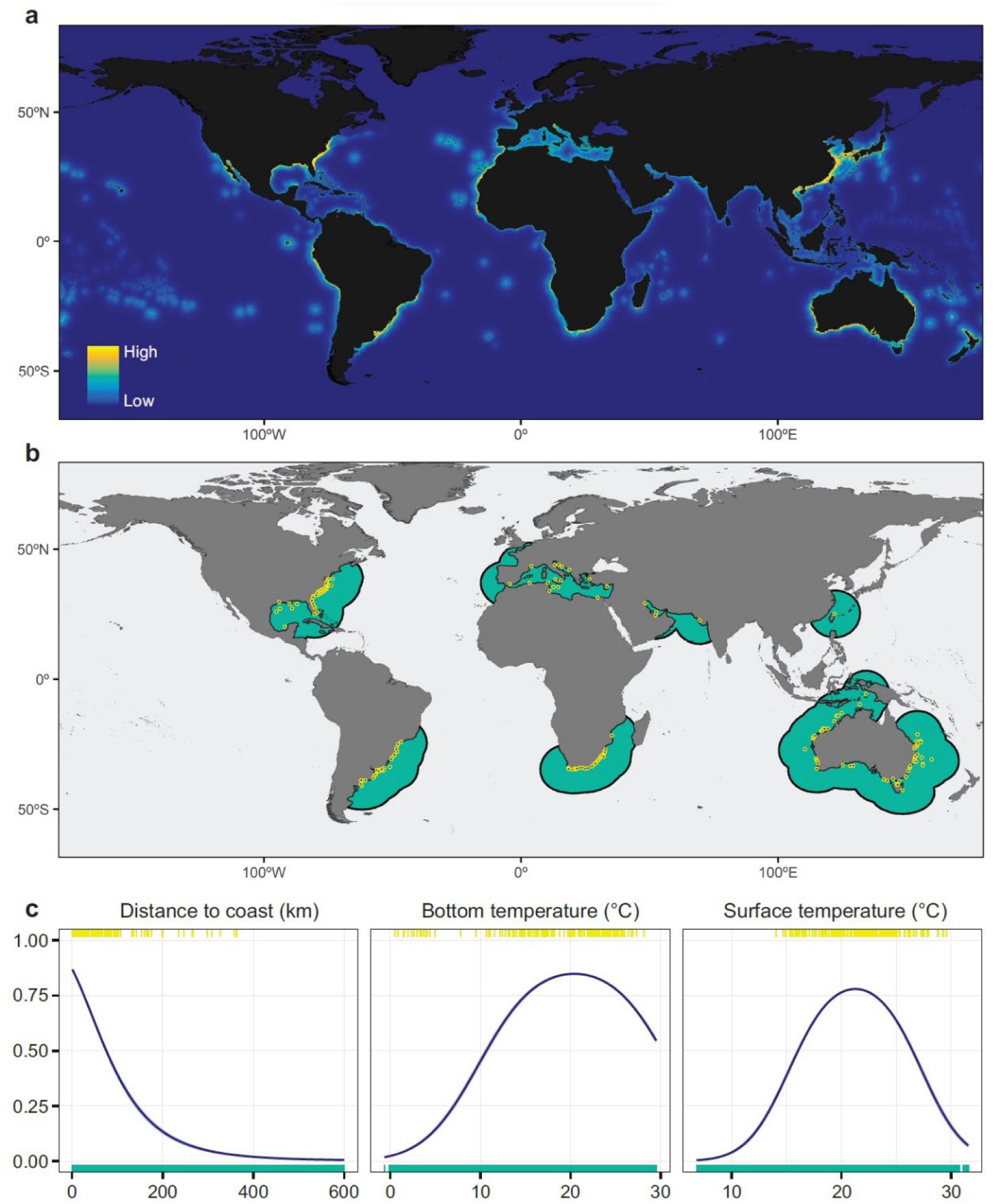
Main model results for the global population of *Carcharias taurus*, including: (**a**) Continuous prediction of habitat suitability representing the potential distribution of the species worldwide, covering both occupied and potentially invadable areas. Areas of strict extrapolation due to model transfer were removed. (**b**) Calibration areas constructed as 1000 km diameter buffer circles around each of the calibration points, represented as circles. (**c**) Response plots (logistic output) of the most influential predictors in the model. The curves illustrate how the prediction varies as the values of each predictor are modified, while keeping the rest of the predictions at their average value. The lines represent the mean response, and the shaded area corresponds to one standard deviation of variability from ten MaxEnt replicates. At the top, predictive values corresponding to calibration points are shown, while at the bottom, values corresponding to 10,000 background locations randomly taken from the calibration area are displayed.

The model predicts that suitable habitat for *C. taurus* is widely distributed in the area of interest, the SWA (Figure 4a). The areas of potential distribution were found in coastal and continental shelf zones, spanning from southern Brazil (∼18°S) to central Argentina (∼41°S) (Figure 4b). The estuarine areas of the Río de la Plata, areas beyond the continental slope, and those south of the parallel 42°S were not suitable for the species. All historical occurrences of *C. taurus* in the region were recorded in areas predicted as suitable for the species (Figure 4a).

**Figure 4.**
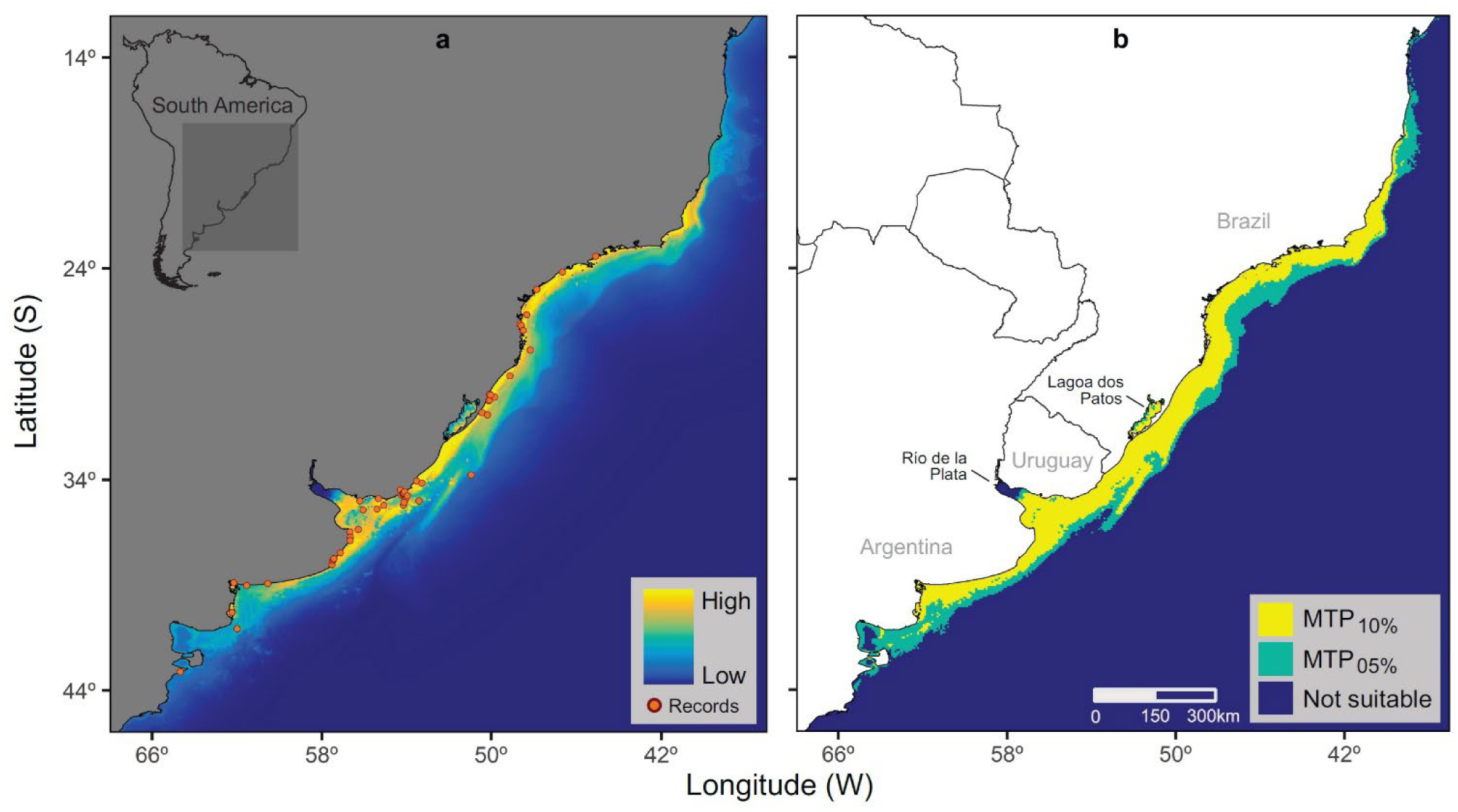
Continuous (**a**) and binary (**b**) potential distribution of the *Carcharias taurus* population in the Southwest Atlantic, based on the indirect estimation of its ecological niche at a global scale. The predictions are not limited to the calibration area but represent the model transfer to the entire region, while areas of strict extrapolation have been removed. Historical occurrences of the species in the region are shown in **a**. The binary prediction was established using minimum training presence (MTP) thresholds with 10% and 5% error on the habitat suitability prediction.

## 4. Discussion

By integrating various occurrence and environmental data in this study, ENMs provided a comprehensive understanding of the habitat requirements and potential distribution of widely distributed mobile marine species populations. This approach is particularly valuable for species like *C. brachyurus* and *C. taurus*, where data may be sparse or unevenly distributed across their regional range. Focusing on habitat use and distribution at the population level, this study provides insights into the broader ecological patterns and processes that govern the distribution of shark species in the SWA. This population-level perspective is essential for understanding the dynamics of species spatial (niche) overlaps, resource use, and ecosystem functioning, offering a holistic view of the species’ ecological roles and requirements.

The identification of potential suitable habitats through ENM is crucial for the development of effective conservation strategies. By pinpointing areas essential for the survival of these species, resource managers can prioritize efforts in habitat protection and implement measures to mitigate human impacts, such as fishing pressures and habitat degradation. Additionally, the modelling approach helps identify potential habitats that have not been extensively surveyed, guiding future research and monitoring resources. This is critical for improving our knowledge of the spatial ecology of species of conservation concern and ensuring that management actions are based on robust scientific evidence. The strategies discussed here are not only applicable to *C. brachyurus* and *C. taurus* but also to other widely distributed marine species facing similar data scarcity issues. By adopting a flexible and integrative approach, combining global datasets with local validation efforts, researchers can overcome the limitations posed by data scarcity and contribute valuable information to marine conservation efforts.

### 4.1 Carcharhinus brachyurus

The global map confirmed the potential distribution in the four broad regions where previous studies have shown that *C. brachyurus* occurs abundantly, including the SWA, Southwest Indian Ocean, Australia, and New Zealand (Smale, 1991; Francis, 1998; Lucifora et al., 2005; Rogers et al., 2013). The results also indicate significant potential distribution in regions where there are only reports in biodiversity repositories, such as the Northeast and Southeast Pacific, or where only a few individuals have been reported in the literature, such as the Northwest Pacific (Choi et al., 1998; Kim et al., 2022) and the Mediterranean Sea (Hemida et al., 2002; Morey & Massuti, 2003; Storai et al., 2007). In addition, the predicted suitable regions of the Northwest Atlantic and the Azores Islands represent invadable areas for the species.

The indirect habitat suitability predictions obtained for the SWA align with the expected characteristics for a species like *C. brachyurus*, which typically inhabits temperate coastal areas. On one hand, the northern limit of its potential distribution coincided with a recent record south of Brazil near 21°S (Cuevas et al., 2022). On the other hand, the southern limit (Figure 5a) coincided with local ecological knowledge from recreational fishermen in coastal areas of northern Patagonia, who report Puerto Rawson (43.3°S) as the historical boundary of the species’ presence (Irigoyen & Trobbiani, 2016; De Wysiecki, 2014). This information gathered from semi-structured interviews, mapping of traditional fishing grounds, and photograph records confirmed the historical presence of the species seasonally in different points of Northern Patagonia (Río Negro Province: Puerto Lobos; Chubut Province: Playa Fracasso, Playa Canto, and Puerto Rawson), with a southward expansion of about 2° of latitude from the previously documented distribution (Irigoyen & Trobbiani, 2016; De Wysiecki, 2014). Indications of water tropicalization in the region could be the cause of a gradual and progressive expansion process of *C. brachyurus* distribution towards higher latitudes (Franco et al., 2020).

**Figure 5.**
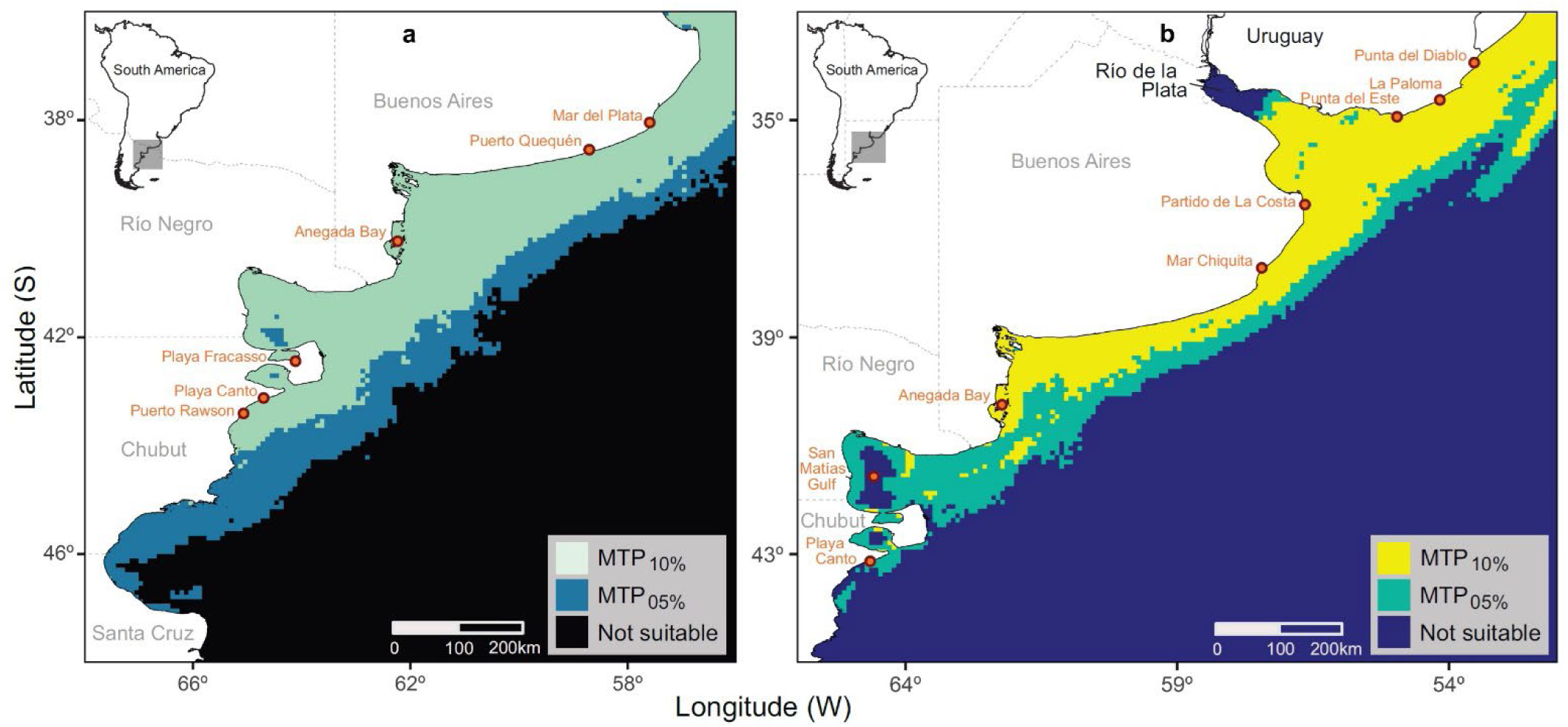
Southern limit of potential distribution of the *Carcharhinus brachyurus* (**a**) and *Carcharias taurus* (**b**) populations in the Southwest Atlantic. The locations discussed in the text are indicated for reference. The binary prediction was established using minimum training presence (MTP) thresholds with 10% and 5% error on the habitat suitability prediction.

Temperature and distance to the coast were the most determining predictors in the habitat use of *C. brachyurus* at the population level, which is consistent with previous studies at smaller spatial scales, both in the SWA and in other regions of the world. Overall, local ecological knowledge and records collected on social media support the frequent capture of the species by recreational fishing in Argentina, which usually occurs during the warm season between October and May in much of the Buenos Aires Province (Irigoyen & Trobbiani, 2016; De Wysiecki, 2014). Similarly, local studies demonstrate a marked seasonality of the species during the months of highest water temperature. For example, off Puerto Quequén (38.6°S) and Mar del Plata (38°S), captures of *C. brachyurus* have been recorded in the weeks prior to December, suggesting that the progressive warming of coastal waters southward favours the use of these habitats by the species (Chiaramonte, 1998a, 1998b; De Wysiecki, 2024). About 350 km south, in the coastal area of Anegada Bay (40.2°S) in northern Patagonia, it was observed that the increase in water temperature during December coincides with the first captures of *C. brachyurus*, extending until April, while the water temperature is unsuitable for the species for the rest of the year (Lucifora et al., 2005). In southern areas, such as Puerto Lobos (42°S), recreational fishing of *C. brachyurus* also begins in December but is only recorded during the summer months (De Wysiecki, 2024). In the SWA, therefore, temperature seems to drive the coastal distribution and habitat use of *C. brachyurus*, whereas the availability of a particular prey is unlikely to drive movement patterns (Lucifora et al., 2009). On the South African coast, water temperature is also key factor, both directly, as a physiological limitation, and indirectly, by affecting the distribution of its main prey in this region, the sardine *Sardinops sagax* (Cliff & Dudley, 1992; Dudley & Cliff, 2010). Furthermore, an increase in detections of tagged individuals of *C. brachyurus* was observed in a gulf off southern Australia and in a bay off northern New Zealand coinciding with the seasonal increase in water temperature (Drew et al., 2019; Kellett, 2021). Therefore, seasonal variation in water temperature appears to be one of the main factors driving the movements and distribution of *C. brachyurus* in coastal environments worldwide.

A subset of historical occurrences of *C. brachyurus* in the SWA was recorded in areas distant from the coast that were predicted as unsuitable for the species. That is, these records occupied environmental combinations that were not representative of the those occupied at a global scale by the species, especially habitats significantly more distant from the coast. This result was striking as it represents a large amount of data for the region (*n* = 134, 32.2%), although its representation at a global scale was very low (<1%). The most plausible explanation for this result is that the subset of records corresponds to captures made exclusively by pelagic longline fishing. In fact, these records come from only two sources of information: 132 records from the Uruguayan pelagic longline fleet (Mas Bervejillo, 2012) and two records from a Brazilian ichthyological collection captured with pelagic longline (Soto & Mincarone, 2004). This fishing gear allowed for the collection of captures of *C. brachyurus* in the water column, in areas far from the coast, where benthic habitats can exceed 500 m in depth. At a global scale, no other datasets replicating these sampling conditions for *C. brachyurus* were found, explaining why its ecological niche was located closer to the coast. This highlights the limitations of ENM, where the accuracy of predictions is intimately linked to the representativeness of available data and their ability to capture the relevant environmental conditions for the species. Therefore, it is likely that suitable habitats for *C. brachyurus* also include neritic areas away from the coast, which the species uses for specific purposes, such as foraging or migration. For instance, if there existed pelagic longline fishing in Argentina, there could be catches of this species. This would alter the current understanding in the region, suggesting that *C. brachyurus* inhabits neritic shelf areas much more frequently than previously thought. In fact, some recent studies suggest that the distribution of *C. brachyurus* extends to neritic shelf habitats where the depth exceeds 100 m (Bradshaw et al., 2018; Kellett, 2021). Consequently, it is necessary to incorporate representative data from the water column to obtain more accurate future estimates of the ecological niche of *C. brachyurus*.

### 4.2 Carcharias taurus

The global map confirmed the potential distribution in the four broad regions where previous studies have shown that *C. taurus* occurs abundantly, including the SWA and Northwest Atlantic, Southwest Indian Ocean, and Australia (Branstetter & Musick, 1994; Lucifora et al., 2002; Pollard et al., 1996; Smale, 2002). The results also indicate significant potential distribution in regions where there are only reports in biodiversity repositories, such as the Northwest Pacific, or where only a few individuals have been reported in the literature, such as the Persian Gulf (Jabado et al., 2013) and the Mediterranean Sea (Bargnesi et al., 2020). In addition, the predicted suitable regions of the East Pacific, Galápagos Islands, Azores Islands, Madagascar, and New Zealand represent invadable areas for the species.

The indirect habitat suitability predictions obtained for the SWA align with the expected characteristics for a species like *C. taurus*, which frequents subtropical and temperate coastal areas. The northern limit of the potential distribution in southern Brazil was located at approximately 18°S, suggesting that the species could use coastal areas farther north than the most extreme documented records to date at nearly 24°S (Motta, 2006). Regarding the southern limit (Figure 5b), the prediction was located near the San Matías Gulf, coinciding with the most extreme historical records of the species (Menni et al., 1981). In this regard, the only capture recorded in the 1990s at Playa Canto (43.1°S), along with the ecological knowledge of recreational fishermen that the presence of *C. taurus* in Chubut is unusual or rare and has not been observed again since the date of the interviews, reinforces the idea that the 41°S parallel effectively marks the southern geographical distribution limit of this species (Irigoyen & Trobbiani, 2016; De Wysiecki, 2024). This finding coincides with the southern limit (∼41°S) of the Australian east population around the Bass Strait and northern Tasmania (Ebert et al., 2021).

As for *C. brachyurus*, temperature and distance to the coast were the most determining predictors in the habitat use of *C. taurus* at the population level, which is supported by previous studies at smaller spatial scales in the SWA. Local ecological knowledge and records collected on social media also suggest that the species is frequently caught by recreational fishing between November and April in much of the Buenos Aires Province, Argentina (Irigoyen & Trobbiani, 2016; De Wysiecki, 2024). During the warm months in Mar Chiquita (37.7°S), significant captures of the species are observed in recreational fishing activities, with the possibility of sighting it associated with specific rocky formations (De Wysiecki, 2024). In the coastal area of Anegada Bay (40.2°S), a strong seasonality in *C. taurus* captures was observed, with an increase in water teemperature between December and April appearing to be a conditioning factor, while the rest of the year the thermal conditions of the habitat would be inadequate for the species (Lucifora et al., 2002). These records support the hypothesis that the increase in water temperature during certain times of the year promotes greater use of these coastal areas by *C. taurus*. In Uruguay, for example, there is a historically understudied artisanal fishery that relies on *C. taurus* to make “bacalao criollo” in Punta del Diablo (34°S) (Beretta et al., 2023), suggesting that coastal catches are common. Additionally, field surveys and local ecological knowledge indicate catches of the species are also frequent, including La Paloma (34.7°S), Punta del Este (34.9°S) and nearby areas (Praderi, 1985; Silveira et al., 2018). However, its capture in artisanal fisheries in areas adjacent to the mouth of the Río de la Plata (i.e. Partido de La Costa), was reported as rare during summer and winter, likely due to highly fluctuating and potentially inadequate estuarine conditions (Jaureguizar et al., 2015). Overall, there is a need for studies on monitoring recreational and artisanal fishing, along with non-invasive methods, to determine other essential habitats for the species, such as nursery and breeding areas. Although studies on *C. taurus* in the SWA are limited, water temperature appears to be the main driver of its distribution and habitat use in coastal areas.

In other parts of the world, water temperature has been shown to be a determining factor in the large-scale coastal distribution of *C. taurus*. On the South African coast, the first captures of neonates and juveniles of the species occur in summer, when water temperature exceeds 18°C, and extend until autumn, when these values decrease (Dicken et al., 2006, 2007). Similarly, adult individuals marked on these coasts moved in response to seasonal and interannual temperature fluctuations in a range of values between 10 and 22°C (Smale et al., 2012). On the east coast of Australia, *C. taurus* exhibits a habitat use pattern mainly associated with water temperatures in the range of 17 to 24°C and typical depths less than 40 m (Otway & Ellis, 2011). Correspondingly, at an aggregation site off the north coast of Western Australia, the species’ habitat use depends largely on water temperature staying below a threshold value of 24°C (Hoschke & Whisson, 2016). In bays on the east coast of the United States, temperatures below 18°C were determinants in the emigration of juveniles from nursery areas (Kneebone et al., 2012, 2014) and in the migration of adults (Teter et al., 2014; Haulsee et al., 2018). At smaller spatial scales, water temperature is also an important variable in the spatial distribution of *C. taurus*, both directly (Kneebone et al., 2012, 2018) and the distribution of relevant prey (Dicken et al., 2007). In summary, seasonal variation in water temperature appears to be one of the main factors conditioning the movements and distribution of *C. taurus* in coastal environments in different parts of the world (e.g., Smale et al., 2015; Dwyer et al., 2023).

### 4.3 Data limitations and ENMs

While the ENM approach utilized in this study offers valuable insights into the habitat use and potential distribution of shark populations in the SWA, it is important to acknowledge the potential impact of incomplete sampling or observation efforts on the accuracy of our findings. As highlighted by Peterson & Soberón (2012), the ecological niche might be underestimated if sampling efforts are incomplete across the study area. The limitations imposed by incomplete sampling efforts can introduce biases into the data used for modelling, leading to gaps in our understanding of species’ ecological requirements and distribution patterns (Hortal et al., 2008). In the case of *C. brachyurus* and *C. taurus*, where data may be sparse or unevenly distributed across their range, these biases could result in an incomplete representation of their ecological niches. This is particularly relevant when considering regions where there are limited reports in biodiversity repositories or where only a few individuals have been documented in the literature. For example, our study identified significant potential distribution areas for both species in regions where historical records are scarce or rely on anecdotal evidence, such as the Northeast and Southeast Pacific for *C. brachyurus*, and the Persian Gulf and the Mediterranean Sea for *C. taurus*. However, the reliability of these predictions may be influenced by the availability and quality of data from these regions, which could be affected by factors such as underreporting, sampling biases, or data accessibility issues. Furthermore, the subset of historical occurrences of *C. brachyurus* recorded in areas distant from the coast, which were predicted as unsuitable for the species, highlights the importance of considering the representativeness of available data in ENM. While these records provide valuable insights into the species’ distribution beyond coastal habitats, they also underscore the need for caution when extrapolating niche predictions based on limited or biased datasets.

In cases where data availability is severely limited or where data quality is questionable, it may be prudent to refrain from attempting ENM altogether. Modelling under such circumstances could lead to misleading or unreliable results, potentially exacerbating rather than mitigating uncertainties in species distribution assessments. Instead, researchers should prioritize efforts to improve data collection and validation processes, leveraging alternative methodologies such as expert consultation or qualitative assessments where quantitative modelling is not feasible. In recognition of the challenges posed by data scarcity, it was determined that attempting to model the direct regional distribution of *C. brachyurus* and *C. taurus* in the SWA was not feasible due to the limited availability of regional data. Consequently, a global modelling approach was implemented to provide a broader understanding of the potential distribution of these species across their entire range. Overall, incorporating representative data from a variety of habitats and sampling conditions is necessary for improving the accuracy and reliability of future estimates of species’ ecological niches.

### 4.4 Conclusion

Addressing the challenges of data scarcity in marine ecological studies requires innovative solutions and adaptive strategies. ENMs are rarely used in marine systems, making this a fairly new approach that holds significant promise for enhancing our understanding of marine biodiversity. While predicting habitat use from ENMs is often a first step in understanding for data-poor regions with severely depleted populations, the strength of ENM is that even without effort data, essential habitats for a species can likely be identified. By leveraging global data, integrating citizen science contributions, and employing advanced modelling techniques, researchers can improve the accuracy and applicability of ENMs. This approach can provide critical insights into the distribution and habitat use of marine species, ultimately supporting more effective conservation and management efforts in data-scarce regions. The application of ENM in this study has enhanced our understanding of the habitat use and potential distribution of *C. brachyurus* and *C. taurus* in the SWA. The findings highlight the importance of specific environmental factors in shaping the distribution patterns of these species and underscore the value of niche modelling as a tool for conservation planning. By predicting broader potential distribution patterns, this study lays a foundation for informed and effective management of shark populations in the SWA. Furthermore, the ENM approach appears to be fit for purpose for area-based conservation initiatives such as ISRA (Hyde et al., 2022).

## Acknowledgements

This study is part of the doctoral thesis of ADW, supported by a scholarship from Consejo Nacional de Investigaciones Científicas y Técnicas (CONICET, Argentina). No directed funds were used for the development of this study. ACM thanks the BlueMarineFoundation for their support. Pesca Nación (Argentina), Instituto Nacional de Investigación y Desarrollo Pesquero (INIDEP, Argentina), Australian Fishery Management Authority (AFMA) and Fisheries New Zealand (FNZ) kindly provided presence data.

## Data availability statement

Open presence records from this study are available in the Supporting Information. The code to perform all analyses can be accessed at https://github.com/Agustindewy/Copper_Stiger_shark_SWA

