## Supplementary material for "Novel use of global occurrence data to indirectly predict suitable habitats for widely distributed marine species in data-scarce regions"

#### Supplement 1

Supplementary results of the present work. Includes one table and seven figures.

**Table S1.1** Scientific publications (peer-reviewed and grey articles) related to the study of *Carcharhinus brachyurus* and *Carcharias taurus* in the Southwest Atlantic consulted during the development of this study. The "General" category includes publications that are not specific to the species of study but were consulted for including several species at once or for having the potential to include them, e.g. listings, keys, landings, etc. The literature review was completed on February 28, 2023. References for each publication are linked in each citation.

| Species | Publications |
|---|---|
| *C. brachyurus* | Chiaramonte (1996), Cuevas *et al.* (2018, 2022), Lucifora *et al.* (2005a, 2009a), Soto (2000) |
| *C. taurus* | Cardoso *et al.* (2011), Cuevas *et al.* (2021), Lucifora *et al.* (2001, 2002, 2003, 2009b), Nisa-Castro-Neto (2013), Tanzola & Sardella (2006), Sadowsky (1969b), Soto (2003) |
| General | Alemany *et al.* (2021), Almerón-Souza *et al.* (2018), Barbini & Cousseau (2015), Barbini *et al.* (2015), Barletta *et al.* (2010), Barreto *et al.* (2017), Bernardo *et al.* (2020), Bornatowski & Abilhoa (2012), Bornatowski *et al.* (2009, 2017, 2018b), Bovcon *et al.* (2013), Caille *et al.* (1995), Cedrola *et al.* (2009b, 2011, 2012), Cervigón & Cousseau (1971), Chiaramonte (1998a, 1998b), Colonello *et al.* (2014), Costa & De Tarso Da Cunha Chaves (2006), Cousseau *et al.* (2019), Crespi-Abril *et al.* (2013), Delpiani G. *et al.* (2020), Delpiani S. M. *et al.* (2020), Domingo *et al.* (2017), Figueroa (2019), Flaquer Da Rocha & Ferraz Dias (2015), Funes *et al.* (2022), Gadig (2001), Galina (2006), García *et al.* (2008), Góngora *et al.* (2009), Haimovici (1998), Hornke (2017), Irigoyen & Trobbiani (2016), Irigoyen *et al.* (2015), Jaureguizar *et al.* (2003, 2004, 2006, 2015, 2016), Klippel *et al.* (2016), Knoff *et al.* (2001), Kotas *et al.* (2017), Laporta *et al.* (2018), Ligrone *et al.* (2014), Llompart *et al.* (2010), Lucifora *et al.* (2012), Marcovecchio *et al.* (1988, 1991), Marín *et al.* (1998, 2020), Mas *et al.* (2014), Menezes (2010), Menezes *et al.* (2003), Menni & García (1985), Menni & Lucifora (2007), Menni *et al.* (1984, 1985, 1986, 2010), Milessi *et al.* (2019), Motta (2006), Nani (1964), Otero *et al.* (1982), Paesch *et al.* (2014), Perier *et al.*, (2011), Praderi (1985), Reyes & Torres-Florez (2009), Ringuelet & Aramburu (1960), Romero *et al.* (2013), Ruibal Núñez *et al.* (2018), Sabadin *et al.* (2020, 2022), Santos *et al.* (2021), Schejter *et al.* (2012), Silveira *et al.* (2018a), Soto & Mincarone (2004), Spier *et al.* (2018), Trobbiani *et al.* (2021), Van Der Molen *et al.* (1998), Velasco *et al.* (2007), Viana *et al.* (2017), Vögler *et al.* (2020), Vooren (1997), Vooren & Klippel (2005) |
| Doctoral Theses (Argentina) | Lucifora, 2003, Llompart, 2011, Molina, 2012, Chiaramonte, 2015, Cuevas, 2016, Sabadin, 2019 |

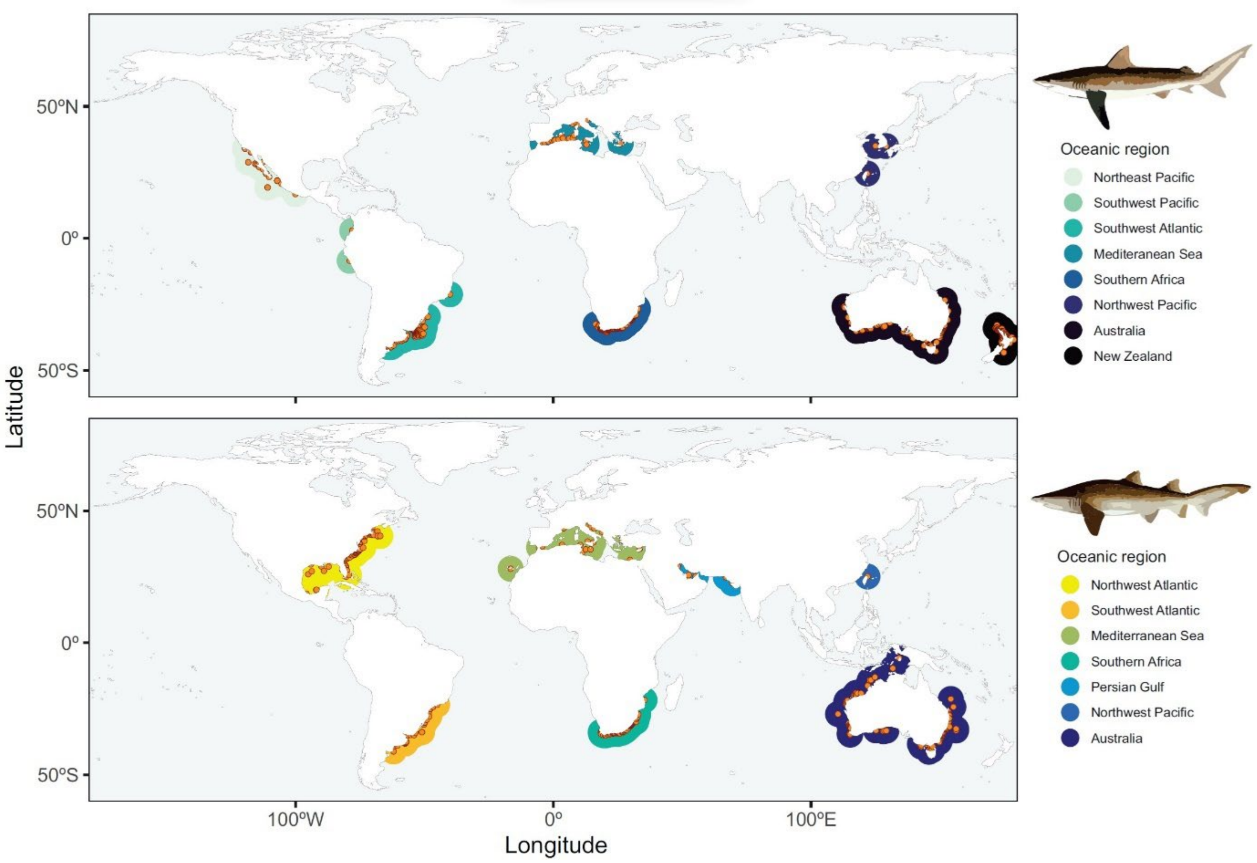


**Figure S1.1** Global distribution of the open-access occurrence records of *Carcharhinus brachyurus* (upper panel, *n* = 11170) and *Carcharias taurus* (lower panel, *n* = 2541) sharks, differentiated by oceanic regions. A 1000 km diameter area around each record (points) was included to facilitate visualization. The data were collected from a variety of freely available sources, including peer-reviewed scientific articles, grey literature, biodiversity repositories, and social media. Metadata for these records can be found in the associated Excel file, and collection details in the Methods section.

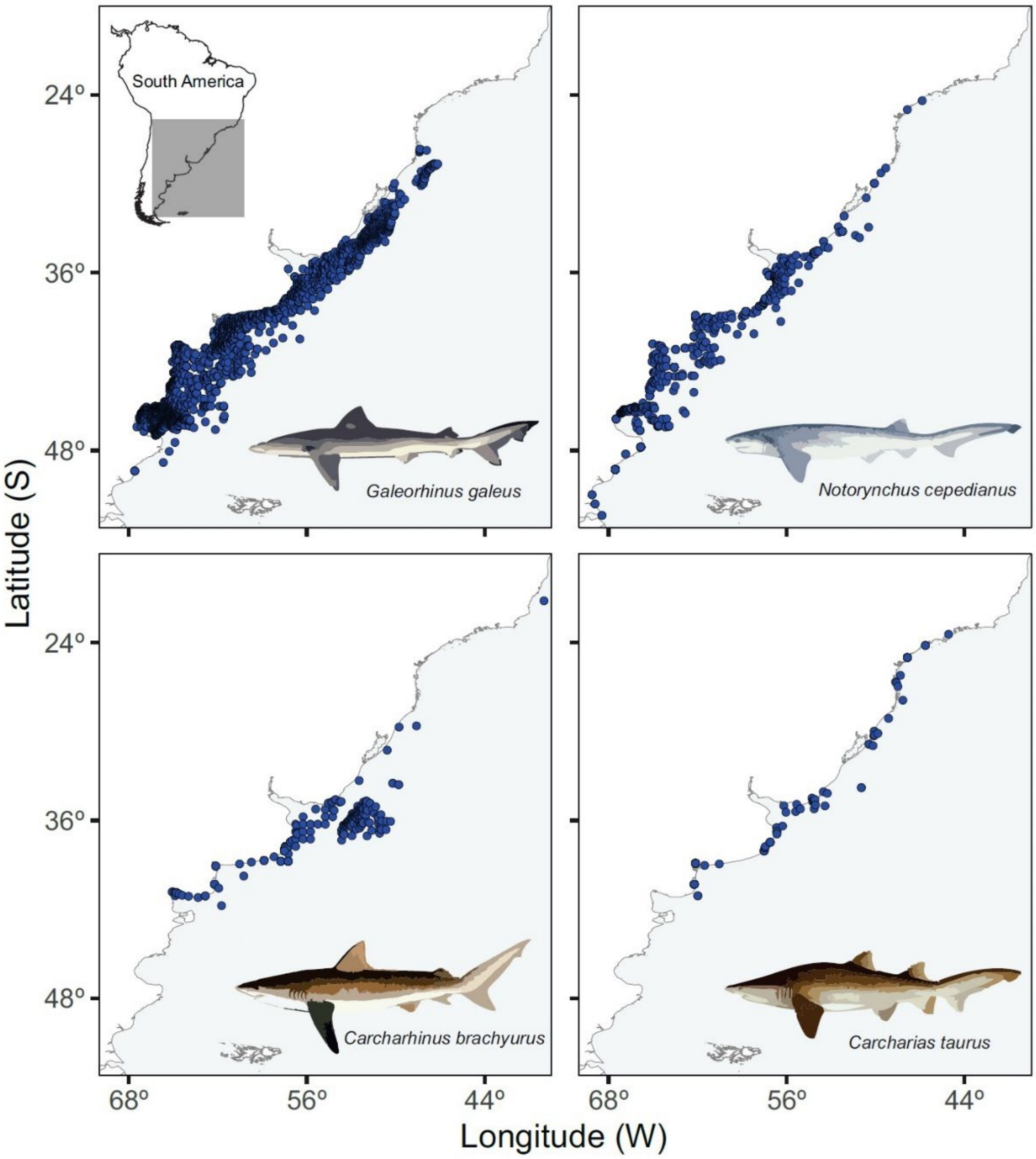


**Figure S1.2** Location of all occurrence records collected for *Notorynchus cepedianus* (*n* = 2591), *Galeorhinus galeus* (*n* = 9611), *Carcharhinus brachyurus* (*n* = 416) and *Carcharias taurus* (*n* = 120) sharks in the Southwest Atlantic. The data were collected from a variety of sources, including peer-reviewed scientific articles, grey literature, scientific surveys, biodiversity repositories, and social media (see Methods section for details). Metadata for these records can be found in the associated Excel file, and collection details in the Methods section.

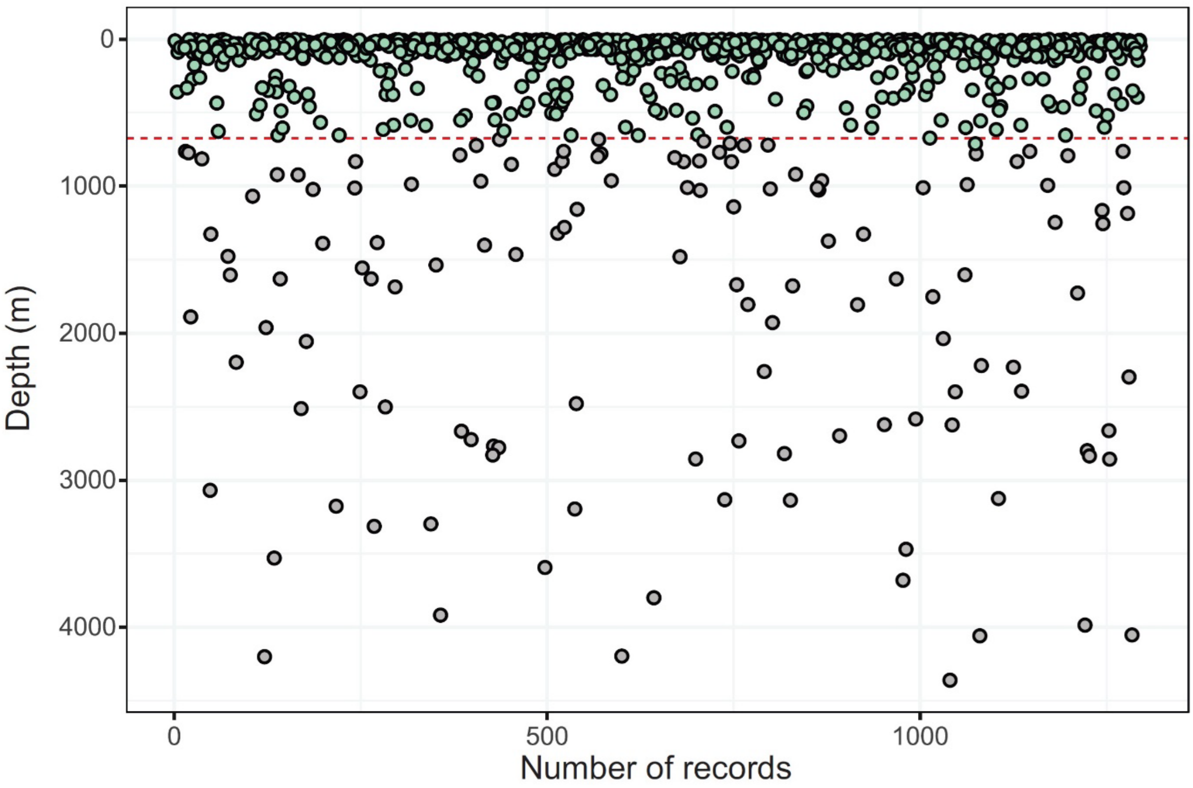


**Figure S1.3** Depth of the records collected for *Carcharhinus brachyurus* estimated using the global bathymetric layer. The dashed line represents the actual maximum depth recorded globally for this species, which is 731 m. Note the unexpected depth values below the dashed line due to discrepancies in the bathymetric layer arising at the margins of the continental shelf. Duplicate captures in space were removed for visualization ease, resulting in a total of 1295 records.

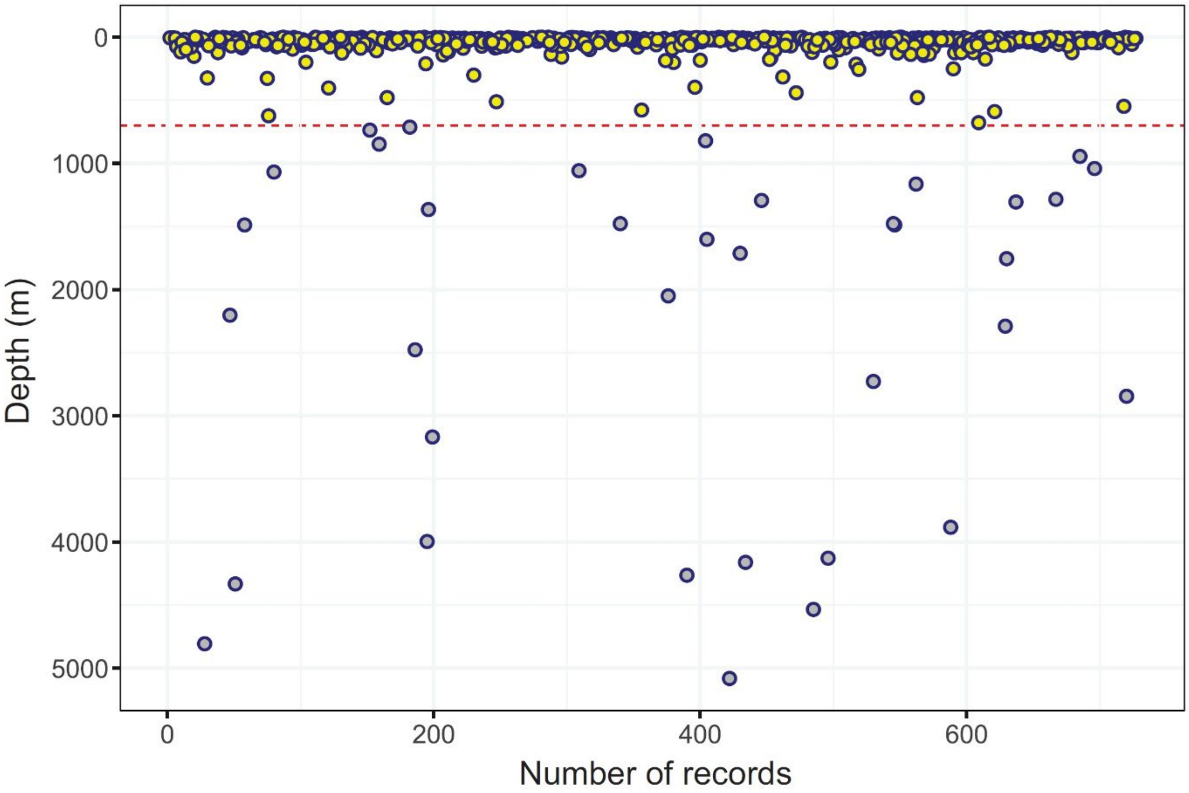


**Figure S1.4** Depth of the records collected for *Carcharias taurus* estimated using the global bathymetric layer. The dashed line represents the actual maximum depth recorded globally for this species, which is 700 m. Note the unexpected depth values below the dashed line due to discrepancies in the bathymetric layer arising at the margins of the continental shelf. Duplicate captures in space were removed for visualization ease, resulting in a total of 727 records.

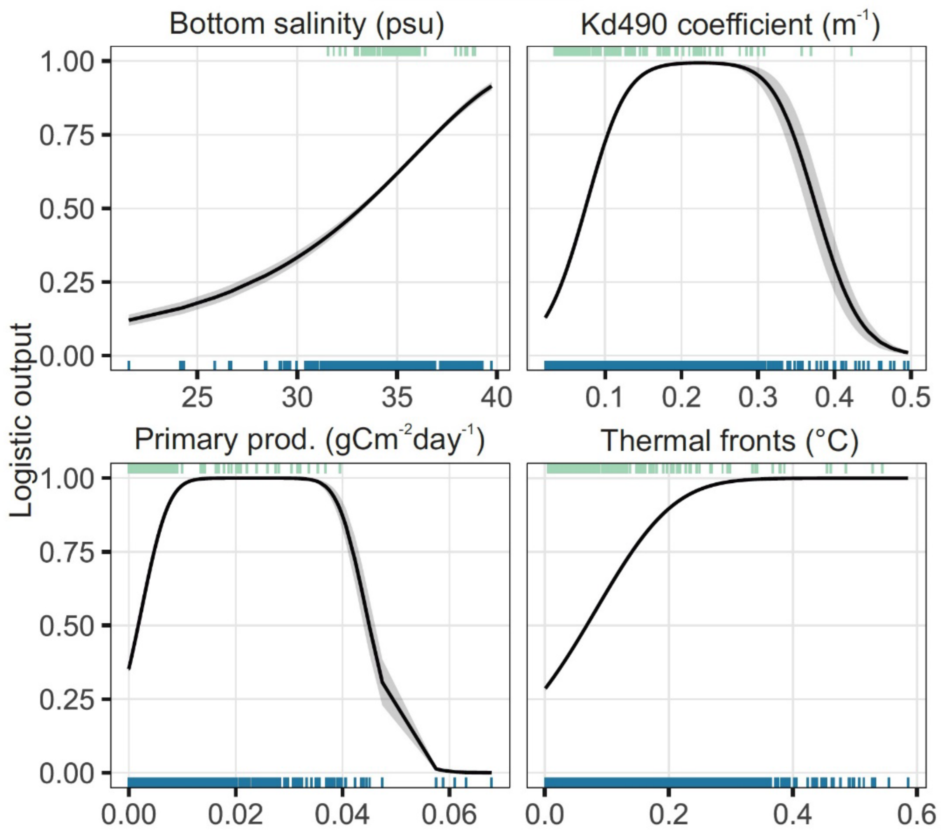


**Figure S1.5** Response plots of marginal predictors for the annual model for the global population of *Carcharhinus brachyurus*. The curves illustrate how the prediction varies as the values of each predictor are modified, while keeping the rest of the predictions at their average value. The lines represent the mean response, and the shaded area corresponds to one standard deviation of variability from ten MaxEnt replicates. At the top, predictive values corresponding to calibration points are shown, while at the bottom, values corresponding to 10,000 background locations randomly taken from the calibration area are displayed.

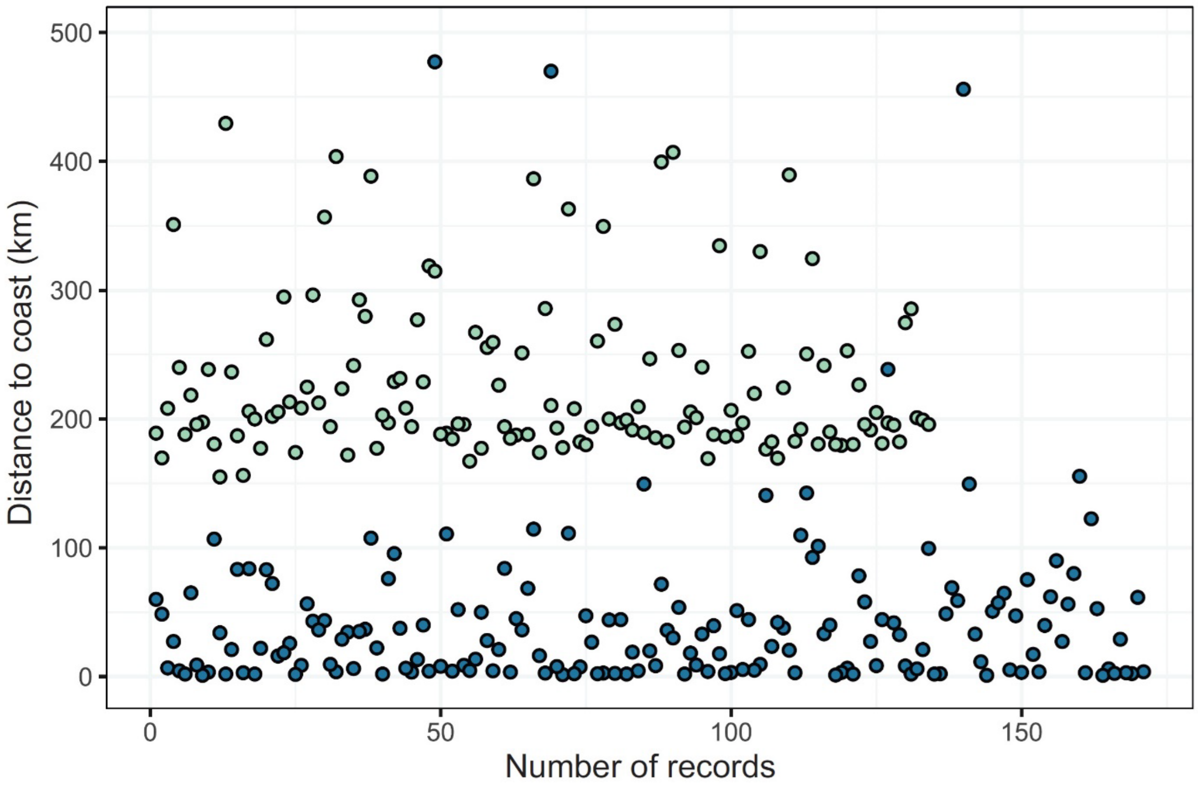


**Figure S1.6** Differences between the distance to the coast values occupied by the calibration points of the global model of *Carcharhinus brachyurus* (*n* = 171, dark circles) and the historical occurrences of the species in the Southwest Atlantic (*n* = 134, light circles) that were found outside the areas predicted as suitable by the model.

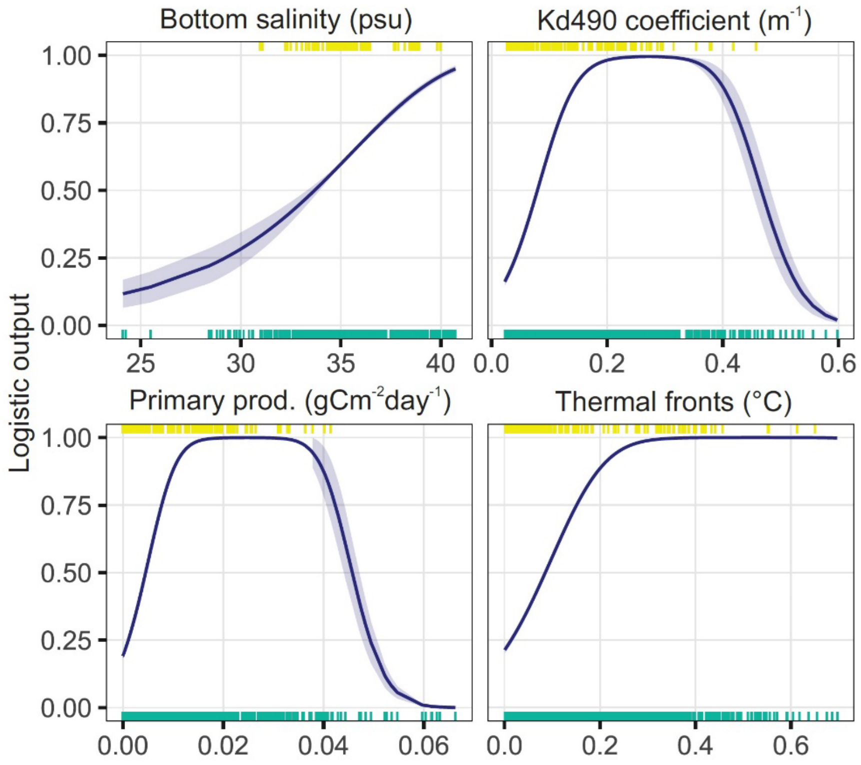


**Figure S1.7** Response plots of marginal predictors for the annual model for the global population of *Carcharias taurus*. The curves illustrate how the prediction varies as the values of each predictor are modified, while keeping the rest of the predictions at their average value. The lines represent the mean response, and the shaded area corresponds to one standard deviation of variability from ten MaxEnt replicates. At the top, predictive values corresponding to calibration points are shown, while at the bottom, values corresponding to 10,000 background locations randomly taken from the calibration area are displayed.

**Supplement 2**

**ODMAP file.**

### Novel use of global occurrence data to indirectly predict suitable habitats for widely distributed marine species in data-scarce regions

#### – ODMAP Protocol –

2024-06-22

---

#### Overview

##### Model objective

Model objective: Mapping and transfer

Target output: continuos and binary habitat suitability

##### Focal Taxon

Focal Taxon: The copper (*Carcharhinus brachyurus*) and sand tiger (*Carcharias taurus*) sharks

##### Location

Location: Global for calibration, worldwide and specifically the Southwest Atlantic for transfer

##### Scale of Analysis

Spatial extent: -180, 180, -90, 90 (xmin, xmax, ymin, ymax)

Spatial resolution: ~0.083º

Temporal extent: from 1810 to 2021

Temporal resolution: single period

Boundary: natural

##### Biodiversity data

Observation type: citizen science, field survey, satellite and acoustic tracking, standardized monitoring data, social media

Response data type: presence-only

##### Predictors

Predictor types: climatic, habitat

##### Hypotheses

Hypotheses: Based on the historical knowledge about copper and sand tiger sharks' distribution worldwide, we hypothesize that the species potential distribution include temperate and subtropical coastal regions of the world.

##### Assumptions

Model assumptions: Both species are in equilibrium with the environment. All relevant predictors are included in the models. Records are representative of the whole distribution of the species globally. Sampling biases and spatial autocorrelation were accounted for. The world's populations of both species exhibit similar habitat use patterns.

##### Algorithms

Modelling techniques: maxent

Model complexity: candidate models with differing complexity (varying feature classes and regularization multipliers) were evaluated.

Model averaging: none.

##### Workflow

Model workflow: automated calibration and evaluation protocol for MaxEnt to compare candidate models of differing complexity (Cobos et al. 2019; Kass et al. 2021). This protocol allows selecting the most significant, best-performing, and simplest models.

##### Software

Software: modelling was performed through the ENMeval and kuenm R packages (Cobos et al. 2019; Kass et al. 2021) that uses the maxent.jar file (version 3.4.1).

Code availability: R scripts available at GitHub (Agustindewy/Copper_Stiger_shark_SWA)

Data availability: most records compiled in this study are made available in Supporting Information, while raw occurrences from Pesca Nación (Argentina), AFMA (Australia) and FNZ (New Zealand) are excluded because they have confidentiality agreements in place. However, all occurrence point data used for calibration are available in Supporting Information.

#### Data

##### Biodiversity data

Taxon names: *Carcharhinus brachyurus* and *Carcharias taurus*

Ecological level: species, populations

Data sources: Catch records in commercial fishing from Pesca Nación (Argentina). Biodiversity data from the Global Biodiversity Information Facility (GBIF, reference below). Biodiversity data from the Ocean Biodiversity Information System (OBIS, reference below).

Catch data available in published and grey literature. Catch data available in social media posts. Catch observer and logbook data from Australian Fisheries Management Authority (AFMA) and Fisheries New Zealand (FNZ).

GBIF.org (27 January 2021) GBIF *Carcharhinus brachyurus* Occurrence Download https://doi.org/10.15468/dl.wdyak3

GBIF.org (27 January 2021) GBIF *Carcharias taurus* Occurrence Download https://doi.org/10.15468/dl.evbzvq

OBIS (Ocean Biodiversity Information System) (2021). Distribution records of Carcharhinus brachyurus (Günther, 1870) and Carcharias taurus Rafinesque, 1810 [Dataset]. Intergovernmental Oceanographic Commission of UNESCO. www.iobis.org (Accessed 27 January 2021)

Sampling design: no sampling design to be reported.

Sample size: the total number of copper and sand tiger shark records retained after data cleaning is 12,704 and 2579.

Clipping: A 1000 km-buffer mask was used for model calibration.

Scaling: records were spatially and environmentally thinned (50 km) to reduce spatial clustering and potential sampling biases.

Cleaning: odd presence data were removed based on Cobos et al. (2018) recommendations. Presences on land were removed.

Background data: 10,000 background records were randomly obtained from calibration areas in each case. Calibration areas corresponded to polygons based on 1000-km buffer areas around spatially and environmentally thinned occurrence records.

#### Data partitioning

Training data: models were generated using a spatial block cross validation procedure based on blockCV R package (Valavi et al. 2019).

Validation data: training data subsets were used to create candidate models and testing data subsets to evaluate them based on significance (partial ROC) and omission rates (10% omission error). Then, model complexity was calculated on models created with the complete set of occurrences (AICc scores). Please refer to Cobos et al. (2019) for further details.

#### Predictor variables

Predictor variables: Distance to the coast (kilometers). Surface temperature (degrees Celcius). Diffuse attenuation coefficient Kd490 (1/meters). Botom temperature (degrees Celcius). Bottom salinity (PSU, practical salinity units). Primary productivity (g C $m^{-2}$ $day^{-1}$, grams of Carbon per square meter a day). Magnitude of thermal fronts (degrees Celcius).

Data sources: the distance to the coast was downloaded at a resolution of ~1 arcmins from the Global Self-consistent, Hierarchical, High-resolution Geography Database (GSHHS, version 2.3.7, URL: http://www.soest.hawaii.edu/pwessel/gshhg/index.html) (Wessel & Smith, 1996). The other variables were downloaded from the global Bio-ORACLE marine dataset (version 2.0) at a resolution of ~5 arcmins (Tyberghein et al. 2012; Assis et al. 2017, URL: https://www.bio-oracle.org/).

Spatial extent: -180, 180, -90, 90 (xmin, xmax, ymin, ymax)

Spatial resolution: distance to coast: ~1 arcmins; others: ~5 arcmins

Coordinate reference system: EPSG: 4326

Temporal extent: 2000-2014

Temporal resolution: single time frame

Data processing: layers were masked by the shoreline polygon and the calibration areas. The distance to the coast variable was resampled to 5 arcmins using the aggregate function in the 'raster' R package.

##### Transfer data

Data sources: the variables were downloaded with a global extent. Spatial resolution, temporal extent and resolution, and data processing were the same as the calibration.

Spatial extent: -180, 180, -90, 90 (xmin, xmax, ymin, ymax)

#### Model

##### Variable pre-selection

Variable pre-selection: none.

##### Multicollinearity

Multicollinearity: multicollinearity between predictors was addressed via the Pearson correlation analysis. Only weakly correlated (-0.7–0.7) predictors were retained.

##### Model settings

maxent: featureSet (lq, lp, lqp), featureRule (We only considered the subset of features that yield the simplest responses, characterised by linear and bell shapes), regularizationMultiplierSet (Sequence from 0.1 to 2 by 0.1 increments, plus 2.5, 3, 4, 5, 7, and 10). Output format (logistic).

Model settings (extrapolation): simple extrapolation to the world.

##### Model estimates

Parameter uncertainty: final model uncertainty was considered from two sources. One, the effect of 10 bootstrap replicates of each best model. Two, the effect of different parametrization of median averaging of best-selected models following protocol in Cobos et al. (2019).

Variable importance: predictor importance was assessed with the jackknife analysis included in the MaxEnt algorithm.

##### Model selection - model averaging - ensembles

Model averaging: none.

##### Analysis and Correction of non-independence

Spatial autocorrelation: no non-independence analyses were carried out.

##### Threshold selection

Threshold selection: binary predictions based on 10th and 5 th Minimum Training Presence (MTP). We considered areas with habitat suitability above the threshold as 'suitable' and those below as 'not suitable'.

#### Assessment

##### Performance statistics

Performance on validation data: most significant (partial receiver operating characteristic scores, partial ROC) and best-performing (omission rate at a threshold E = 10%; i.e., false-negative rate) models among candidate models were narrowed down based on the Akaike's information criterion corrected for ample size (AICc). A ΔAICc value of ≤2 was chosen as a criterion to select the best few models from among each candidate model set. Please refer to Cobos et al. (2019) for further details.

##### Plausibility check

Response shapes: we used partial dependence plots to check the ecological plausibility of fitted species-environment relationships.

#### Prediction

##### Prediction output

Prediction unit: Predictions of habitat suitability expressed as continuous and binary scales.

Post-processing: No post-processing.

##### Uncertainty quantification

Algorithmic uncertainty: Range uncertainty around 10 bootstrap replicates in each model calibration.

Input data uncertainty: Not applicable.

Scenario uncertainty: Not applicable.

Novel environments: Novel environments were assessed thought the Mobility Oriented Parity (MOP, Owens et al. 2013) analysis to identify and rule out areas of strict extrapolation worldwide. The MOP metric compares multivariate environmental distances between the selected background and the global region to which models are transferred. Values of zero indicate strict extrapolation.
